# Optimizing Soybean Breeding: High-Throughput Phenotyping Technologies for Stink Bug Resistance and High Yields

**DOI:** 10.1101/2025.05.19.654360

**Authors:** Maiara de Oliveira, Alexandre Hild Aono, Patricia Braga, Adriano Abreu Moreira, Fernanda Smaniotto Campion, Felipe Augusto Krause, Fernando Henrique Iost Filho, Juliano de Bastos Pazini, Pedro Takao Yamamoto, José Baldin Pinheiro

## Abstract

The stink bug complex is a major agricultural pest for soybean crops, significantly reducing productivity. Genetic resistance is the most effective control strategy, but its quantitative nature and labor-intensive phenotyping make its implementation in breeding programs challenging. This study explored high-throughput phenotyping (HTP) using unmanned aerial vehicles (UAVs) equipped with RGB cameras to evaluate a soybean population and identify stink bug resistance by correlating image-derived features and machine learning (ML) models. Using an alpha-lattice design with three replications, we phenotyped 304 soybean lines over two seasons under natural stink bug infestations. We manually evaluated five traits associated with stink bug resistance and correlated them with color, texture, and histogram features from aerial images. Three ML models—AdaBoost, SVM, and MLP— were tested to predict these traits. VIs, especially the Visible Atmospherically Resistant Index at the first percentile (VARI_P25) and texture-based indices at 45° and 135°, effectively predicted traits in stressed environments, particularly during flights near maturation. While ML models showed good predictive ability for yield, healthy seed weight, and maturity, they were less effective for stink bug resistance. Increasing the number of UAV flights modestly improved predictive accuracy, though predicting traits across different seasons remained challenging. Despite this, indices like VARI_25P were valuable for screening and excluding less promising genotypes, optimizing breeding program resources. This pioneering work offers valuable insights and highlights the need for further research to optimize resistance selection, promising significant advances in soybean breeding for stink bug resistance.

## Introduction

Soybean (*Glycine max*) is one of the most widely cultivated oilseed crops in the world due to its versatility, being used in the production of oil, meal, biodiesel, and proteins for animal and human nutrition (Hartman et al., 2011). In the 2023/24 season, global soybean production totaled 396.9 million tons (USDA, 2024). Currently, soybean is the most economically important crop in Brazil, which stands out as the largest global producer, achieving an estimated production of 154.6 million tons with an average yield of 3.5 t/ha (CONAB, 2024). Nowadays, soybean is cultivated in almost all regions of Brazil, favoring the emergence of various insect pests and pathogens that cause significant losses in grain yield (GY) and quality (Boerma and Walker, 2005).

Among the main factors compromising soybean productivity in Brazil, stink bugs are particularly detrimental, causing losses of up to 125 kg/ha when considering one stink bug per linear meter (Guedes et al., 2012). This insect feeds directly on soybean pods, affecting the grains and causing losses not only in yield but also in the physiological and phytosanitary quality of the seeds (Bortolotto et al., 2015). Additionally, stink bug’s feeding process causes soybean delayed maturity, which hinders the determination of the harvest time and mechanical harvesting due to leaf retention and green stems (Panizzi and Hirose, 1995; Rossetto et al., 1995).

One of the most efficient control strategies is the use of resistant cultivars, which can be integrated with chemical control. However, stink bug resistance represents a polygenic and quantitative trait, challenging its implementation in breeding programs (Godoi and Pinheiro, 2009). Plant breeding programs targeting quantitative traits require a large and diverse population, which needs to be evaluated in a feasible and precise way. In this sense, phenotyping is one of the main bottlenecks of breeding, as it typically relies on manual and visual measurements, which are time-consuming and imprecise. This limitation restricts the number of lines that can be tested (Bhat and Yu, 2021).

Developing high throughput phenotyping (HTP) methodologies has enhanced the phenotypic evaluation of genotypes, optimizing diverse breeding programs (L. Li et al., 2014). HTP methodologies, particularly utilizing autonomous platforms like unmanned aerial vehicles (UAVs) coupled with cameras, have emerged as valuable tools in assisting breeders in selecting and developing commercial cultivars. Furthermore, the use of images facilitates non-destructive evaluation of numerous traits across a large number of lines at various growth stages and locations simultaneously (Furbank and Tester, 2011; Reynolds et al., 2019).

The incorporation of image-based phenotyping in soybean breeding has been prominent in predicting many important traits, including maturity (Trevisan et al., 2020; Yu et al., 2016), biomass (Moreira et al., 2021; Sakurai et al., 2022), drought resistance (J. Zhou et al., 2020), and yield estimation (Maimaitijiang et al., 2020; J. Zhou et al., 2022). However, the use of HTP for quantifying and identifying insect injury in soybeans is still in its early development.

Marston et al. (2020) demonstrated the effectiveness of using drone-based multispectral imagery and machine learning (ML) models to identify stress caused by soybean aphids, noting a reduction in near-infrared reflectance as aphid populations increased. However, in contrast, Iost-Filho et al. (2022) explored hyperspectral imagery in controlled environments, combined with machine learning (ML) models, and found that infestation by stink bugs, specifically *Euschistus heros* and *Diceraeus melacanthus*, did not influence reflectance patterns in infested or uninfested soybean plants. Thus, applying these technologies in field conditions for the selection of resistant genotypes still requires further research to increase the efficiency and accuracy of the selection process (Goggin et al., 2015).

The present study aims to explore the potential of High-Throughput Phenotyping (HTP) using an Unmanned Aerial Vehicle (UAV) equipped with an RGB camera applied to a population composed of 290 soybean lines. The lines were evaluated over two years using an alpha-lattice design under natural infestation by stink bugs. By contrasting different image descriptors and their association with classical phenotyping measurements, our study provides insights into best practices for soybean image-based phenotyping in evaluating resistance to the stink bug complex. Furthermore, we assessed the efficiency of ML algorithms in predicting several important traits based on image measurements, providing an end-to-end workflow with high potential to assist soybean breeding.

## Materials and Methods

### Plant Material and Field Experiment

For the conduct of this study, we utilized a breeding population consisting of 290 soybean lines, part of the germplasm bank of the Laboratory of Genetic Diversity and Breeding at Luiz de Queiroz College of Agriculture/University of São Paulo (ESALQ/USP). These lines resulted from crosses among the commercial cultivars BRS-133, CD-215, Conquista, Dowling, IAC-100, and Pintado, which were recombined with the plant introduction (PI) genotypes: PI 200487 (Kinoshita), PI 471904 (Orba), PI 200526 (Shiranui), and PI 459025 (Bing Nan). Progenies from these crosses were subsequently recombined with nine populations that exhibited genetic variability for stink bug resistance. These genotypes underwent two selection cycles focusing on resistance to the soybean stink bug complex, in addition to one selection cycle for high GY.

Phenotypic evaluations were performed during the 2019/2020 and 2020/2021 growing seasons in the municipality of Piracicaba/SP (Brazil), at the Anhumas Experimental Genetics Station (22°50’00’ S, 48°05’00’ W, altitude of 460 m). In both seasons, the 290 lines, along with 14 commercial checks recommended for the region and/or possess resistance to stink bugs, were evaluated using a 16×19 alpha-lattice experimental design with three replicates. Experimental plots consisted of two rows, each 4 meters in length, spaced 0.5 meters apart, and with a seeding density of 15 seeds per meter. The experiments were conducted following the recommended agronomic management for soybean cultivation, except not performing chemical control of insects, aiming to ensure the presence of stink bugs in the experimental area, enabling the evaluation of the lines’ resistance to this insect.

To monitor the population and verify the presence of stink bugs in the experimental area, beat cloths were employed from flowering to full maturation. Beat cloths were used weekly throughout the entire reproductive cycle of the plant, covering the period of stink bug occurrence and attack. Evaluations involved the use of a beat cloth along a 1-meter crop row, with random beating across the entire experiment. Adults, nymphs (third, fourth, and fifth instars), and the total population (adults and nymphs) of soybean stink bugs were counted.

### Manually Measured Agronomic Traits

The evaluation of soybean genotypes involved the manual measurement of five key agronomic traits. This encompassed the assessment of the number of days to maturity (NDM), which comprehended the duration from planting until 95% of pods achieved maturity, thereby indicating the genotype’s cycle. Leaf retention (LR) was appraised at maturity using a visual scale ranging from 1 to 5, where a score of 1 denoted plants undergoing normal senescence, while a score of 5 indicated plants retaining multiple stems and green leaves, rendering harvest impractical. GY was determined post-harvest, representing the total mass of seeds produced in the plot, corrected for 13% grain moisture, and converted to kilograms per hectare (kg.ha^-1^). Healthy seed weight (HSW) was determined as the weight of seeds devoid of stink bug damage, evaluated after harvest and grain processing (kg.ha^-1^). This measure was obtained during seed processing, where they undergo separation through a spiral mechanism, where shriveled, green, and malformed grains are separated by the action of gravity and centrifugation (DA ROCHA et al., 2014). Tolerance (TOL) was calculated to assess the genotype’s resilience to stink bug injury, quantified through an index obtained by the following formula:

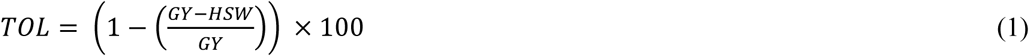

### UAV Data Collection

Aerial images were collected by a DJI Phantom 4 Pro V2.0 drone (DJI, Shenzhen, China) equipped with a 20-megapixel RGB camera capturing images in the visible spectrum. Flight planning was conducted using the FieldAgent digital platform (Sentera, Minnesota, USA) (Fig. 1a). The camera captured sequential images of the area through progressive scanning.

**Fig. 1.**
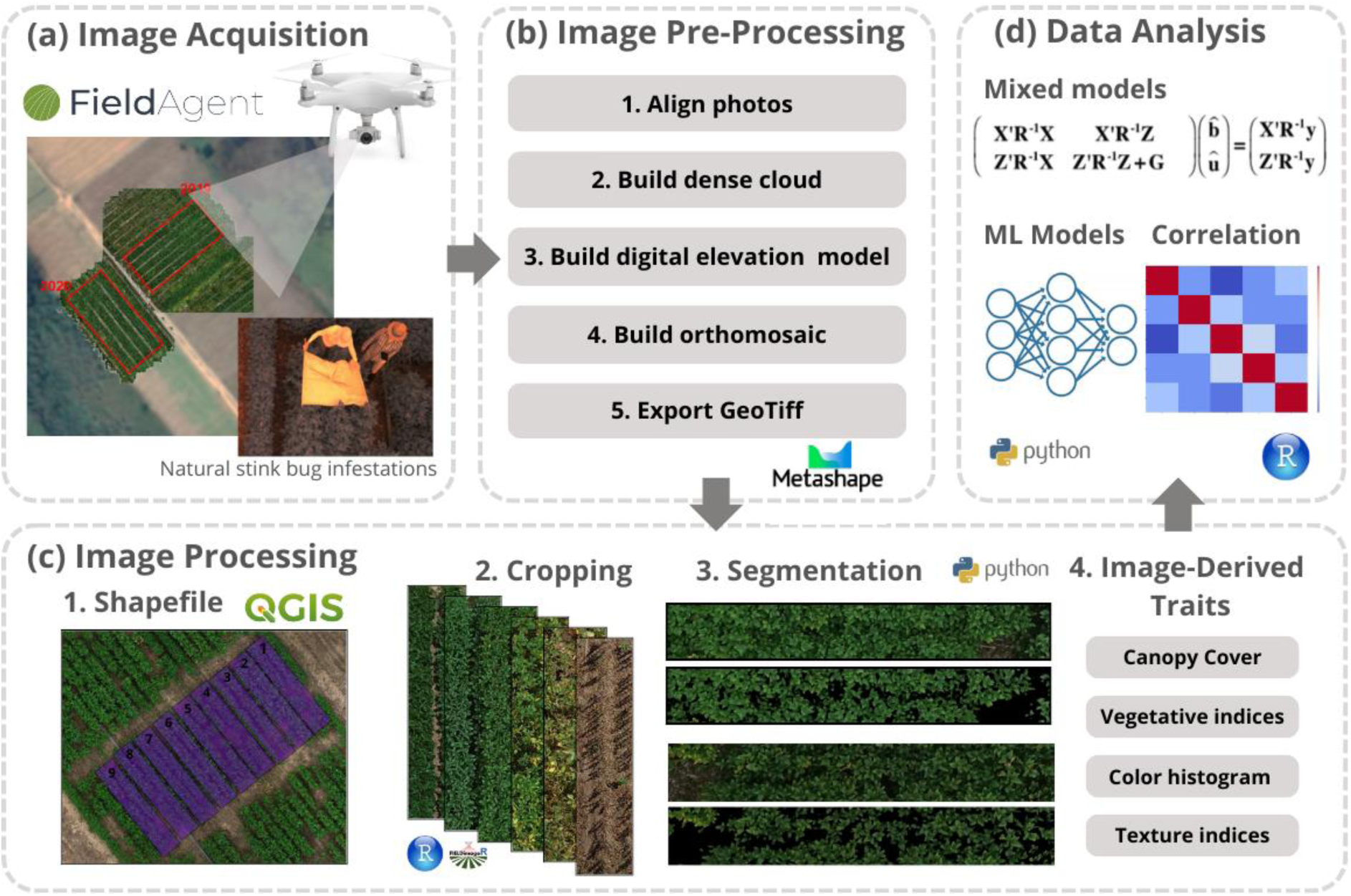
Flowchart of procedures performed, from acquisition to data analysis.

During the crop season of 2019/2020, six flights were conducted, while in the 2020/2021 season, this number was expanded to ten different dates. Detailed information on the days after planting (DAP) when each flight was conducted can be found in Figure 2. All flights were executed at the height of 20 meters, with a ground sampling distance (GSD) of 1.8 centimeters. An 80% level of both frontal and lateral overlap was maintained between the acquired images. Additionally, flights were conducted on sunny, cloud-free days, within the time frame of 10 a.m. to 2 p.m., aiming to optimize lighting conditions and minimize environmental variabilities.

**Fig. 2.**
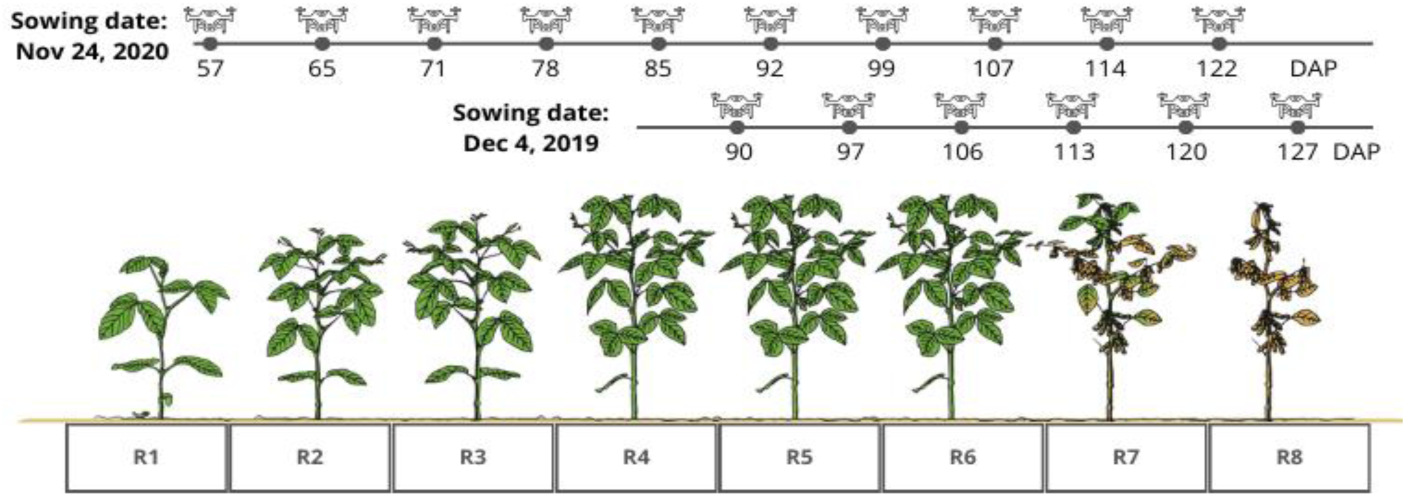
Representation of soybean development stages, indicating the days after planting (DAP) at which drone flights were conducted in the two growing seasons.

### Image Processing

After acquiring the aerial images, individual orthomosaics were created for each field (Fig. 1b) using Agisoft MetaShape software (Agisoft LLC, St. Petersburg, Russia), following the standard protocol provided by the software for orthomosaic generation. Subsequently, the orthomosaics were imported into QGIS version 3.02 software (QGIS Development Team, 2018), where shapefiles were delineated, defining the boundaries of each experimental plot (Fig. 1c.1). To mitigate potential interferences resulting from leaf overlap between adjacent plots, a negative buffer of 0.10 m was applied. Each plot was identified with an ID based on the experimental design. The FieldImageR 0.6.0 package (Matias et al., 2020) implemented in R was used to crop each experimental area simultaneously, employing the shapefile as the plot boundary (Fig. 1c.2).

After individualizing and cropping each plot, background removal was performed, including shadows, soil, and vegetative residues, using the methodology proposed by Yuan et al. (2019), which employs a plant segmentation method using vegetation indices to maximize segmentation accuracy. All the steps were implemented using Python v3 (Van Rossum and Drake, 2009) and the OpenCV library (Pisarevsky, 2008). First, all the plot images were rotated to a 90-degree orientation to calculate the vegetation indices of modified excess green (MExG) and the color index of vegetation extraction (CIVE). By subtracting these indices (MExG - CIVE), we scaled the resulting image into a 0-255 range and used this matrix to create a segmentation mask with Otsu’s thresholding technique (Otsu, 1979) (Fig. 1c.4).

### Image-Derived Traits

For each segmented image corresponding to an orthomosaic at different flight dates, we calculated different vegetation indices and features describing their physical properties, including color, gradient, and texture. We employed the Python v3 programming language (Van Rossum and Drake, 2009) with the libraries OpenCV (Pisarevsky, 2008) and scikit-image (Van Der Walt et al., 2014).

Initially, the Canopy Cover Index (CCI) was computed, representing the percentage of remaining pixels after image segmentation. Additionally, six VIs were calculated (Table 1), including the triangular greenness index (TGI), the green leaf index (GLI), the normalized green-red difference index (NGRDI), the red-green-blue index (RGBVI), and the visible atmospherically resistant index (VARI). Based on the images resulting from each index calculation, we summarized the value distribution into different descriptive statistics: mean, standard deviation (std), skewness (skew), kurtosis, and 25, 50, and 75 percentiles, denoted as P25, P50, and P75, respectively.

**Table 1.** Summary of vegetation indices (VI) used in this study. Each VI was calculated using an RGB image, where the image channels are red (R), green (G), and blue (B).

| Vegetation Index | Name | Formula | References |
| --- | --- | --- | --- |
| **TGI** | Triangular greenness index | $G - 0.39 \times R - 0.61 \times B$ (2) | (Hunt et al., 2005) |
| **GLI** | Green leaf index | $\frac{(2 \times G - R - B)}{(2 \times G + R + B)}$ (3) | (Bassine et al., 2019) |
| **NGRDI** | Normalized green-red difference index | $\frac{G - R}{G + R}$ (4) | (Hunt et al., 2005) |
| **RGBVI** | Red-green-blue index | $\frac{G^2 - R \times B}{G^2 + R \times B}$ (5) | (Bendig et al., 2015) |
| **VARI** | Visible atmospherically resistant index | $\frac{G - R}{G + R - B}$ (6) | (Gitelson et al., 2002) |

To reflect different wavelength values, we created a gray-level histogram with 64 bins, considering each bin quantification as a different attribute. Texture measures were calculated using gray level co-occurrence matrices (GLCMs) to describe repetitive patterns in the image, varying in size and providing information about roughness and regularity. For each image, we constructed four different GLCMs corresponding to angles of 0, 45, 90, and 135 degrees. These matrices were summarized into six different features based in Texture Indices (TIs): energy, correlation, angular second moment (ASM), dissimilarity, and contrast (Haralick et al., 1973).

Therefore, the complete characterization of each image was based on 124 features: 1 feature measuring CC, 7 descriptive statistics for each of the 5 vegetation indices (35 features), 64 features corresponding to color histogram quantifications (Hist_1 to Hist_64), and 6 features for each of the 4 GLCMs (24 features). Associations between these features and the manually measured traits were performed with Pearson correlations in software R version 4.3.2 (R Core Team, 2023).

### Statistical Analysis of Phenotypic Data

The phenotypic data from all trials were analyzed in R (R Core Team, 2023)using the ASReml-R package version 4.3.2 (Butler et al., 2018). Variance components and genetic parameters were estimated using the restricted maximum likelihood (REML) method (Patterson and Thompson, 1971), while genetic values were predicted using the best linear unbiased prediction (BLUP) method (Henderson, 1982). The significance of random effects was assessed through likelihood ratio tests (LRTs).

For the five manually measured traits in the study and each of the six indices obtained from images, statistical analyses were conducted for each year, considering each flight individually. The analyses followed the model:

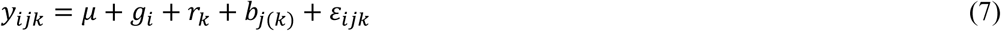

where, *y*_*ijk*_ represents the phenotype observed for the *i*-th genotype, within the *j*-th block, across the *k*-th replicate; *μ* the overall mean; *g*_*i*_ the random effect associated with the *i*-th genotype 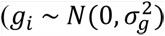 with 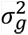 representing the genetic variance); *r_k_* the fixed effect of the *k*-th replicate; *b*_*j*(*k*)_ the random effect of the *j*-th block within the *k*-th repetition 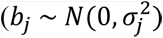 with 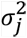representing the block variance); and *ε*_*ijk*_ the model residual, assuming that the residuals are independent and normally distributed, i.e., *ε*_*ijk*_ ∼ *N*(0, *σ*^2^), where *σ*^2^ is the residual variance.

Heritability in the broad sense (ℎ^2^) was estimated using the method proposed by Piepho and Mohring (2007):

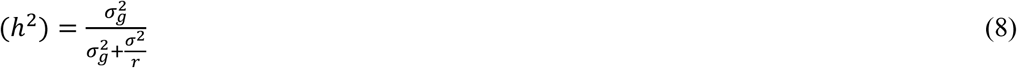

where 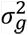 and *σ*^2^ are the genotypic and error variances, respectively, and *r* is the number of replicates. The correlation between the true and predicted random effects was employed as a metric to evaluate the accuracy:

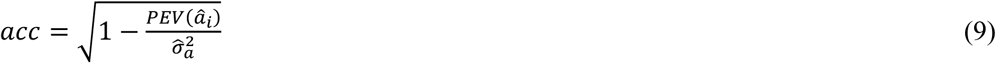

where *PEV*(*â*_*i*_) is the prediction error variance of the predicted random effect and 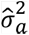 is the estimated variance of the random effect. Using the BLUPs of each genotype, Pearson’s correlations were performed between the traits.

### Machine Learning Methods

The scikit-learn library (Van Der Walt et al., 2014), developed in Python v3 programming language (Van Rossum and Drake, 2009), was employed to implement the ML -based prediction models. We utilized the features calculated from images collected via UAV as independent variables and the five visual characteristics as dependent variables. Each visual characteristic was associated with the UAV-collected indices according to the flight date. For the GY, HSW, TOL and NDM characteristics, regression models were applied, while for LR, a classification model was adopted. Subsequently, six distinct forecasting scenarios were tested. Two scenarios considered all flights conducted in each season (2019/2020 and 2020/2021), while two others focused exclusively on the last three flights of each season. Additionally, two scenarios were designed to test cross-year prediction: the first utilized data from 2019/2020 to predict the phenological characteristics of 2020/2021 based solely on flights near the maturation period, and the second incorporated all flights.

Three machine learning methods were explored in this study: support vector machine for regression (SVR) (Vapnik, 2000), multilayer perceptron (MLP) neural network (Pal and Mitra, 1992), and adaptive boosting (AdaBoost) (Freund and Schapire, 1997). Additionally, we applied a decision tree to investigate which characteristics obtained through images are more important for predicting the manually measured agronomic traits, estimating the feature importance of each image feature using the Gini index.

The performance of the models was evaluated through a 10-fold cross-validation scenario. For the regression models, the Pearson correlation coefficient (*R*) and mean squared error (*MSE*) were used as evaluation criteria. For the classification models, accuracy and precision metrics were employed to assess the model’s ability to classify the test data correctly. These metrics were calculated by comparing the predicted and actual values in the training and test sets.

## Results

### Agronomic Trait Evaluations

To obtain cultivars resistant to stink bugs, the first step is to identify the most promising genotypes. This process faces significant challenges due to the difficulty of phenotyping, as it is a quantitative trait. The use of advanced techniques such as high-throughput phenotyping (HTP) can improve and accelerate the results. However, the implementation of these techniques requires the development of specific protocols for each situation. Furthermore, field evaluation of stink bug resistance is particularly challenging as it depends on the natural infestation of the insects, which can be highly variable between seasons.

To monitor stink bug infestation in the experimental area and assess selection pressure, weekly samplings were conducted using beat cloths throughout the entire reproductive period. It was observed that the natural stink bug infestation remained within the recommended limits for selection in both agricultural seasons studied. As illustrated in Figures 3 and 4, the weekly incidence of stink bugs in the area over the two seasons of the experiment is depicted. Additionally, it can be observed that the dates of the UAV flights coincided with the periods of highest stink bug incidence in the area.

**Fig 3.**
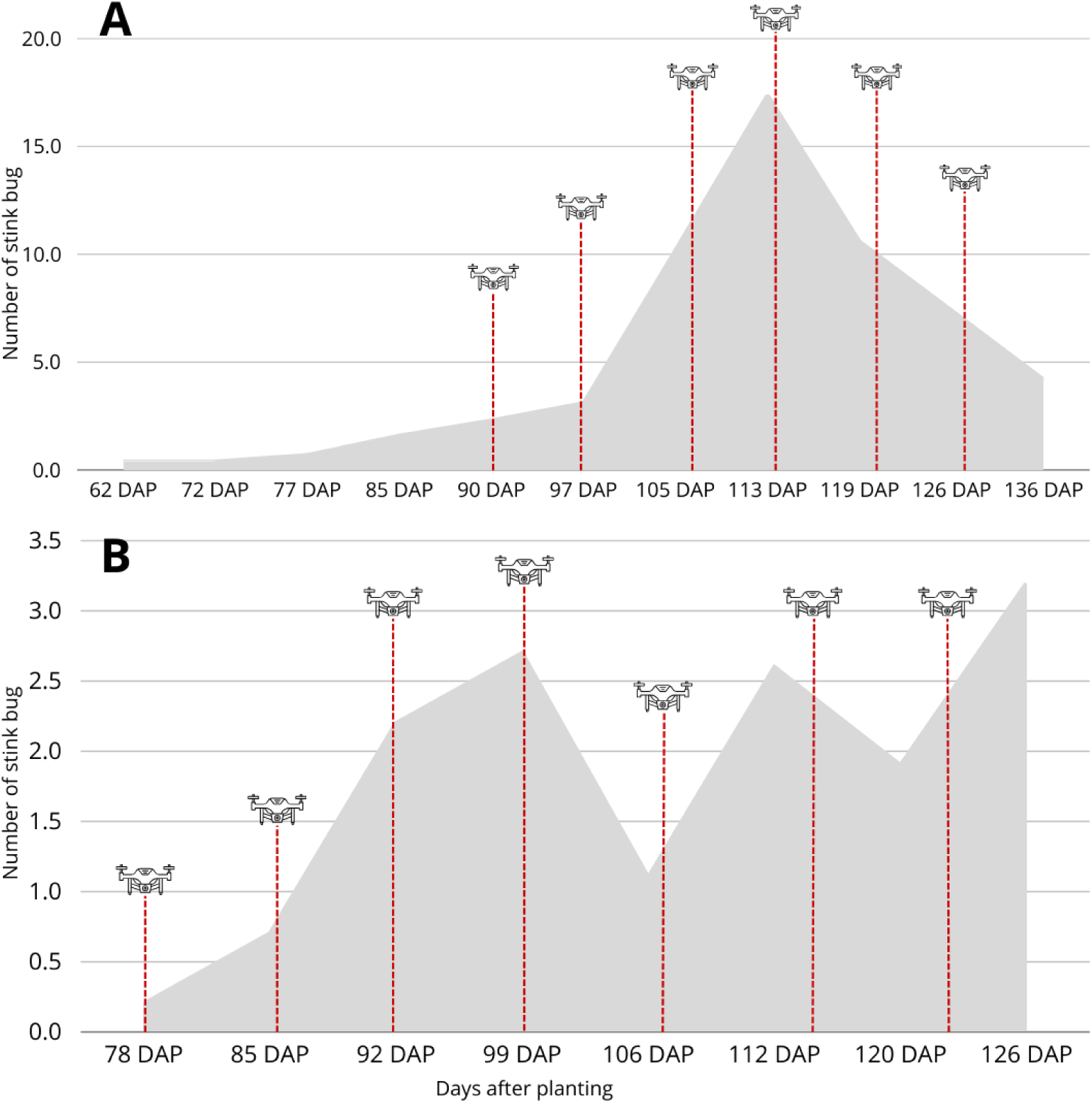
Average occurrence of stink bugs during the reproductive period of soybean in the 2019/2020 (Fig. 3A) and 2020/2021 (Fig. 3B) seasons, obtained through beat cloth sampling. The x-axis represents the number of days after planting (DAP) when the beat cloth sampling was conducted, and the y-axis shows the average number of stink bugs and nymphs counted in the area. The red dotted lines indicate the days when drone flights were performed.

**Fig. 4.**
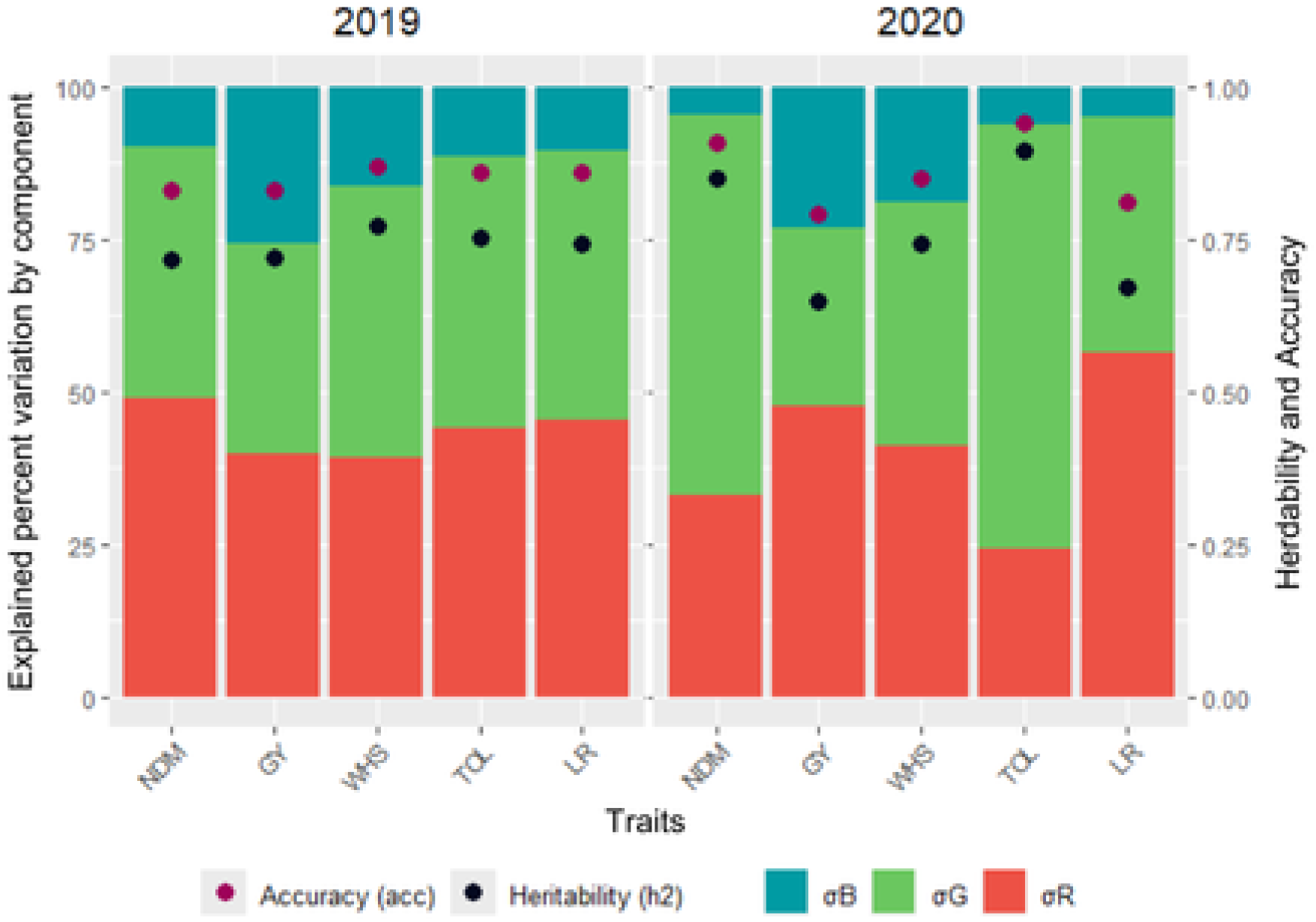
Explained Percent of variation by variance components, heritability, and accuracy for grain yield (GY), hundred-seed weight (HSW), tolerance (TOL), number of days to maturity (NDM), and leaf retention (LR) in two growing seasons.

The 2019/2020 season exhibited a significantly higher infestation, reaching an average of 18 stink bugs per beat cloth at the peak of infestation, 100 days after planting (DAP). In contrast, the subsequent 2020/2021 season had a lower infestation, with a peak of only 3 stink bugs per beat cloth at 120 DAP, along with other smaller peaks recorded at 80 and 100 DAP, averaging 2.5 stink bugs. This variation in infestation can directly impact the identification of resistant genotypes. The lower infestation observed in the 2020/2021 season may make differentiation less evident, as the reduced pest pressure may not fully reveal the resistance capabilities of the genotypes. However, in the 2019/2020 season, with a higher infestation, the more resistant genotypes may stand out more clearly. On the other hand, the infestation may be so high that even the most resistant genotypes, especially those showing tolerance, may not be effective due to the high selection pressure.

To better understand the response of soybean genotypes under natural infestation of stink bugs, it is essential to analyze the variance components that influence the main agronomic traits. The analysis of these components allows for identifying the proportion of total variation attributable to genetic and environmental factors, as well as estimating heritability and measurement accuracy. Figure 4 shows the percentage of variation explained by each model component, along with the heritability (h²) and accuracy (acc) estimates for five manually measured agronomic traits: LR, NDM, GY, TOL, and WHS in each growing season. The variance components considered in the tested phenotypic model were genotype and block. The significance of these components was assessed using the LRT test, which revealed that both components are statistically significant at the 0.01 level, indicating that genetic variations and block variations contribute to the total variation observed in the phenotypic traits.

Genetic variance (σG) accounted for a significant portion of the phenotypic variation across both seasons, demonstrating the strong contribution of genetic factors to trait variation. In 2019, σG accounted for a significant portion of the phenotypic variation, ranging from 34% for GY to 44% for WHS, TOL, and LR, explained between 34% and 44% of the total phenotypic variation. In 2020, while the general trend was consistent, genetic variance for some traits, particularly NDM and TOL, remained high, but was comparatively lower for other traits such as GY and WHS when compared to the 2019 season.

Regarding the significance of the models, both years demonstrated robust estimates of heritability (h²), further confirming the genetic control over the traits studied. In 2019, h² ranged from 0.72 for NDM and GY to 0.77 for WHS, and in 2020, the estimates ranged from 0.65 to 0.90, maintaining the high influence of genetics across the seasons. Prediction accuracy (acc) estimates also reflected this pattern of reliability, ranging from 0.83 to 0.87 in 2019 and slightly higher in 2020, with a range from 0.79 to 0.94, particularly for GY and TOL. These results underline the consistency and reliability of the phenotypic predictions over the two seasons.

In order to evaluate how the manually measured traits are correlated with each other, for a better comprehension of genetic interactions. In Figure 5, we can observe the genetic correlation between the manually measured traits analyzed, with all of them showing significant correlations varying between the two studied seasons. Notably, there is a strong correlation between GY and HSW, the latter often used in the selection of more productive and stink bug-resistant genotypes. This correlation was higher in the 2020 season, reaching a value of 0.87, compared to 0.53 in the 2019 season. However, in the 2019 season, the selection pressure from the presence of stink bugs was considerably high, which may complicate the selection of resistant genotypes. In contrast, in the 2020 season, with lower insect pressure, selection based on HSW may have prioritized more productive genotypes over more resistant ones. This highlights the importance of adjusting stink bug control levels for a more effective selection of soybean genotypes.

**Fig. 5.**
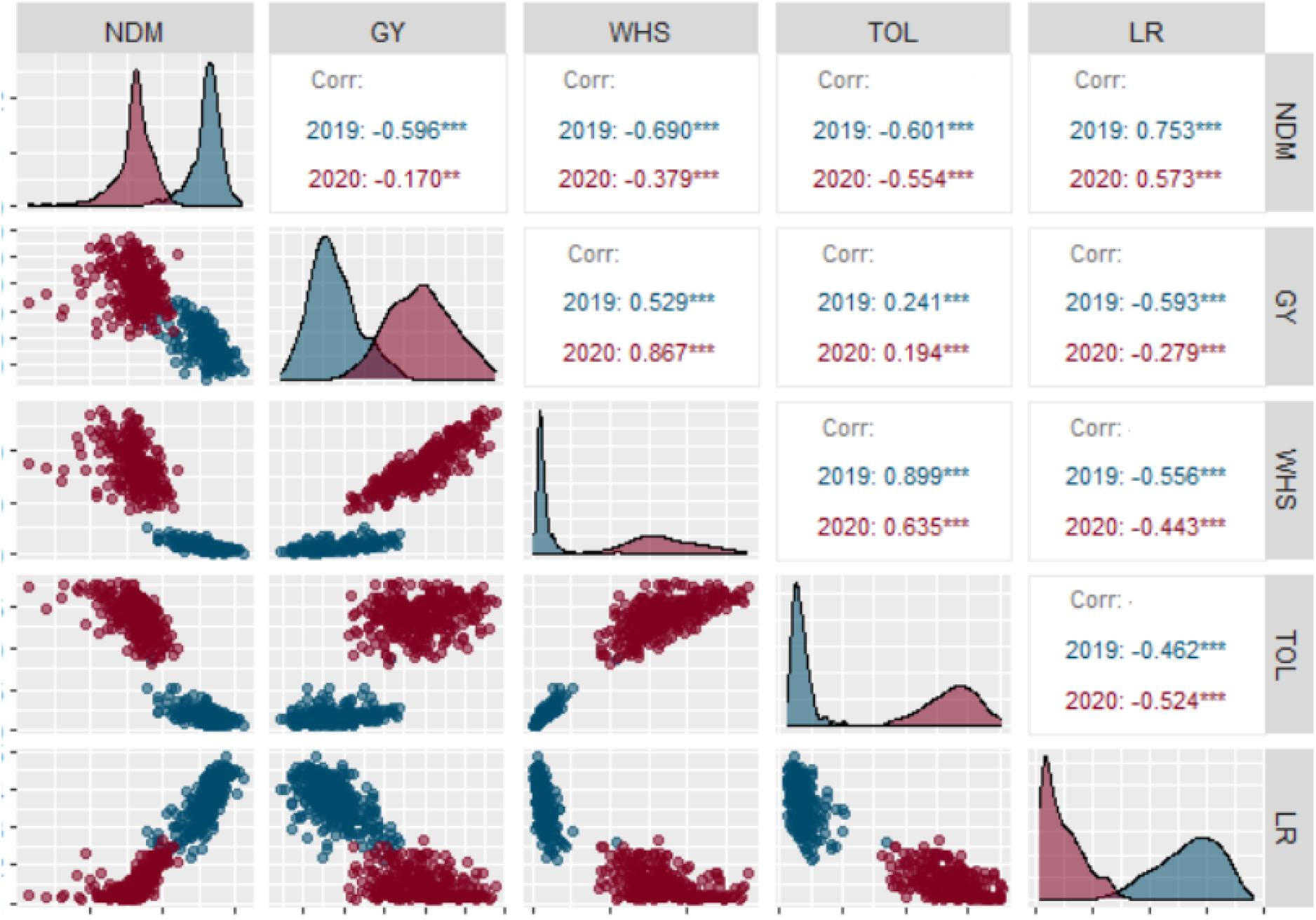
Genetic correlation between the visual characteristics evaluated: grain yield (GY), hundred-seed weight (HSW), tolerance (TOL), number of days to maturity (NDM), and leaf retention (LR) in two growing seasons, 2019/2020 (blue) and 2020/2021 (red).

### Image-Derived Traits

In the 2020 season, we conducted a greater number of flights, starting at 57 days after planting (DAP) and continuing weekly until 122 DAP, totaling 10 flight dates. In contrast, in the 2019 season, only 6 flights were conducted, occurring between 90 DAP and 127 DAP. After the pre-processing and processing stages of the captured images, we obtained 124 indices for each flight date, which included color indices, texture indices, and gradient histograms. To optimize the analysis, we filtered the data based on the correlations of each index with the manually measured traits for each flight. Considering the large number of indices evaluated, we decided to select only the most informative ones for subsequent analyses. Therefore, we selected the indices that showed correlations greater than 0.6, resulting in a filtered set of 38 features.

Among the 38 selected features, 19 were related to vegetative indices (VIs) with various descriptive statistics. Given the high correlation among these indices (Supplementary Fig. 1), we prioritized those with the highest correlations across most flights. This resulted in the selection of four key VIs at the 25th percentile (VARI_P25, RGBVI_P25, NGRDI_P25, and GLI_P25). For the TIs, all those with a correlation greater than 0.6 in any flight were subjected to Principal Component Analysis (PCA), using the first principal component for subsequent analyses (Supplementary Fig. 2). In total, 18 TIs were selected and included in the PCA: ASM, Dissimilarity, Energy, and Homogeneity at angles of 0°, 45°, 90°, and 135°, as well as Correlation at angles of 45° and 135°. These TIs are complementary, as each captures different aspects of the structural and textural variation of the vegetation. Therefore, instead of selecting only a few TIs, we decided to perform a PCA to integrate all the textural information into a single representative index. Additionally, the CC index was included in the selection due to its high correlation with the evaluated traits. In total, six final indices were used (Vari_P25, RGBVI_P25, NGRDI_P25, GLI_P25, CC, and PCA), which were subsequently evaluated in the correlation analyses with manually measured agronomic traits. As the correlations between the manually measured traits and the color histogram quantifications did not exceed the threshold of 0.6, none of them were selected for further analysis.

After selecting the indices, we proceeded with the integration of these indices into mixed models to understand how the comprehension of different characteristics contributes to overall plant performance and facilitates the identification of genotypes with desirable traits. As shown in Fig. 6, the heatmaps illustrate the correlation patterns between image-based indices and DAP across two growing seasons. Correlations between different flight times are stronger for closer periods, while flights that are farther apart in time show weaker correlations. This is evident in the clusters, where closer flight times show significant correlations, whereas more distant flights present correlations close to zero. Given this pattern, each flight time was analyzed individually to understand how different growth stages influence the evaluated traits. By considering each flight time as a distinct dependent variable, we obtain a more precise and detailed analysis, which allows us to identify the specific moment in the crop cycle that is most suitable for performing flights to assess the traits of interest. This approach helps in selecting the ideal time for monitoring each agronomic trait, optimizing the evaluation of genotype performance throughout the different phenological stages.

**Fig 6.**
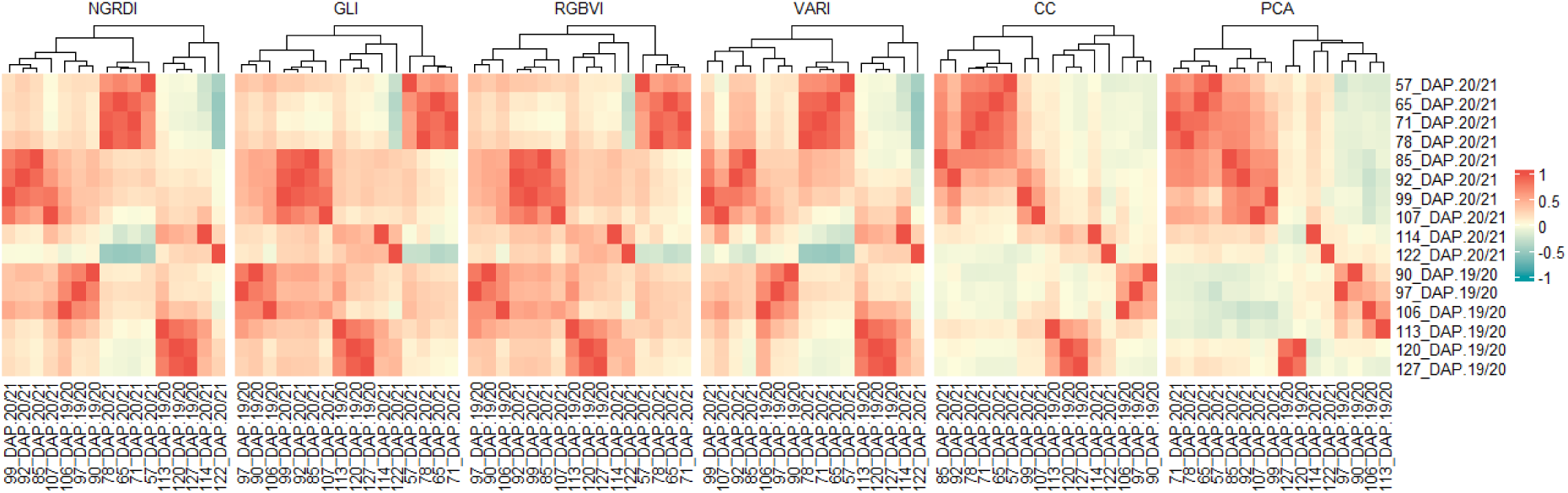
Heatmaps showing the correlation patterns between image-based indices (NGRDI, GLI, RGBVI, VARI, CC, and PCA) and flight times at different days after planting (DAP) during two growing seasons (2019/2020 and 2020/2021).

The decomposition of variance components and the genetic parameters of the six traits obtained through images for each flight are detailed in Supplementary Fig. 3 and 4. It is observed that, across all flight dates, the traits CC and PCA, showed the lowest h² estimates for both seasons. In 2019, flights conducted at 90 and 97 days after planting (grain filling period) presented the lowest h² estimates for all traits. Additionally, the VIs demonstrated similarity in h², with the Vari_P25 at 106 DAP standing out with a h² of 0.76. Overall, the acc estimates ranged from moderate to high, though some flight periods showed lower values. There was considerable variation in acc between different flight periods, reflecting the influence of specific conditions during each period on the accuracy of the predictions. In 2020, the h² for flights conducted at 57, 65, and 122 DAP was generally lower than for other flights. Again, the VI Vari_P25 stood out with the highest variability estimate at 107 DAP (0.77). Acc followed a similar pattern to the previous season. The analysis of variance components revealed that the percentage of genetic variance explaining the observed variability was significant for all traits, as demonstrated by the LRT test, except for PCA at 97 days in the 2019 season. These results suggest the existence of genetic variability that can be exploited, and along with the high h² and acc estimates, indicate that some indices may be important for breeding programs, providing reliable information for the selection of promising genotypes.

Building upon these findings, it is crucial to explore the relationships between the selected image-derived indices and the manually measured traits. Understanding these correlations not only validates the utility of the indices but also enhances the ability to predict agronomically important traits using UAV-captured data, enabling the selection of superior individuals through these indices when a high correlation is found. Fig. 7 present the genetic correlations between indices obtained from UAV-captured aerial images and manually measured traits of soybean at different days after planting (DAP). Strong correlations were observed for all manually measured traits in both seasons, highlighting that certain flight dates are more informative, indicating strong associations with agronomically important manually measured traits and in the evaluation of soybean resistance to stink bugs.

**Fig. 7:**
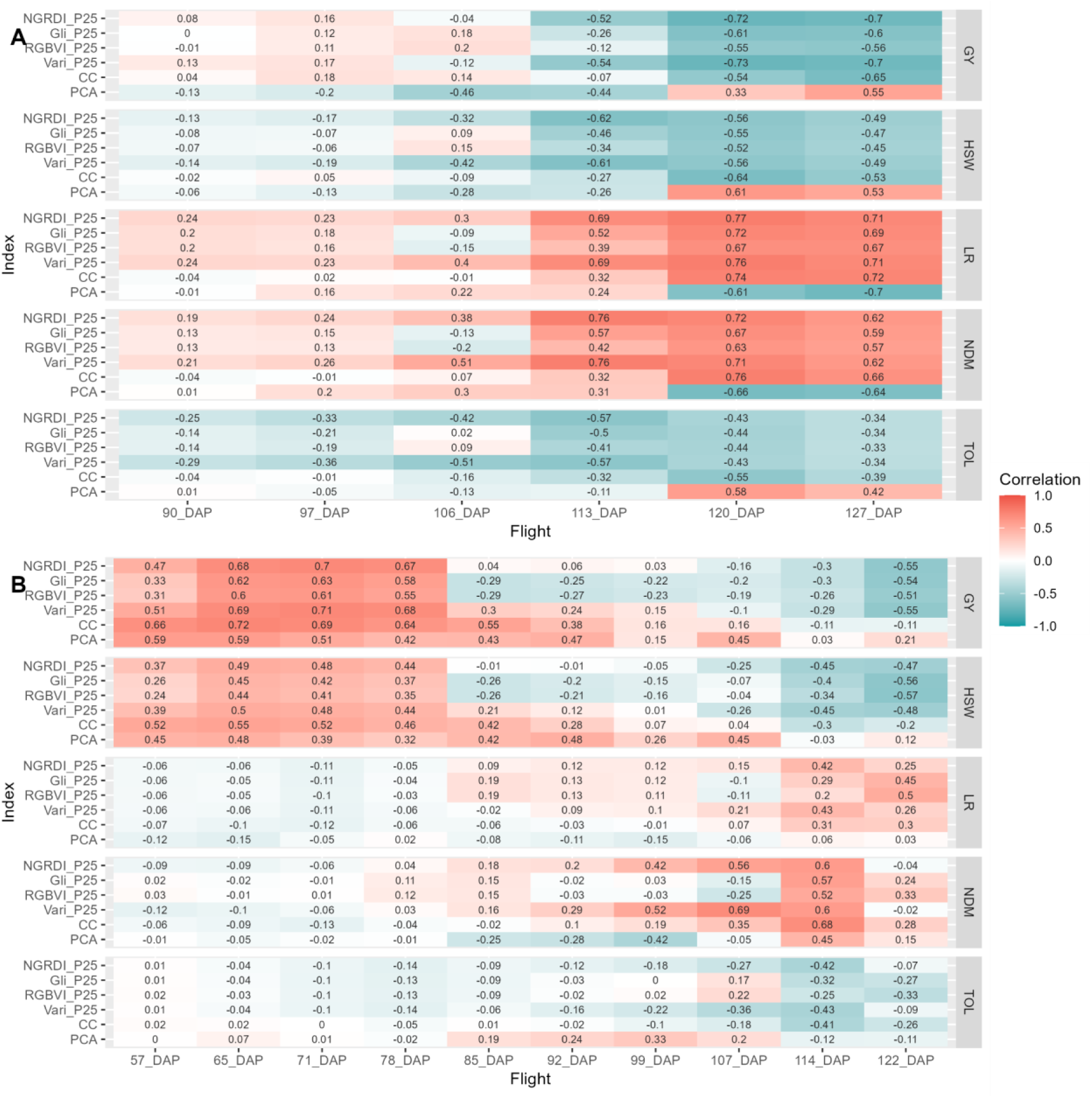
Genetic correlations between image-derived indices and manually measured traits across different flights for the 2019/2020 (Fig 7A) and 2020/2021 (Fig. 7B) growing seasons. Correlations are displayed for various indices (NGRDI _P25, GLI_P25, RGBVI_P25, Vari_P25, CC, PCA) and traits (GY, HSW, LR, NDM, TOL) at multiple DAP (days after planting).

The strongest correlations were found for LR and NDM in the 2019 season, during flights close to harvest and maturity, reaching 0.76. This same pattern was observed in the subsequent season, but with slightly lower correlation magnitudes. For GY and HSW, strong correlations were also found in the 2020 season during the early flights, from 57 to 78 DAP. Additionally, in both seasons, low-magnitude correlations were found between flights occurring from 85 to 107 days for these traits. Moreover, GY and HSW strongly correlated with various indices in the later flights. For TOL, although the correlations were not as strong as with the other traits, in the 2019 season, where stink bug infestation was more pronounced, significant correlations above −0.5 and 0.5 were observed during flights close to maturity with the CC and PCA indices.

The VIs derived from RGB bands exhibit similar patterns among themselves, capturing similar information about vegetation. Additionally, the CC, which is calculated based on soil and leaf area segmentation, shows patterns similar to those of the vegetation indices based on RGB bands. When analyzing the correlations between these indices and the evaluated visual traits, consistent patterns are observed. In the early flights, these indices and the CC show positive correlations with GY, HSW, and TOL, which turn negative in later flights. This shift is due to changes in the growth stages of the plants, where early vigor and biomass positively influence yield and tolerance traits. However, as the plants approach maturity, biomass begins to decrease due to leaf loss, altering these relationships. On the other hand, for LR and NDM, the correlation patterns are opposite. This occurs because plants exhibiting symptoms of leaf retention and delayed maturity tend to have greater biomass at the end of the cycle, as their leaves take longer to fall. Finally, based on texture measures, the PCA index presents distinct patterns from the other vegetation indices. Its response is generally opposite to that of the other indices, suggesting that it captures different information about vegetation structure and spatial distribution.

In summary, these analyses revealed consistent and informative patterns, highlighting that VIs were more effective, with correlations and heritabilities superior to TIs. In particular, the VARI_P25 index stood out by showing high correlations with several important traits in both seasons, in addition to demonstrating consistency across multiple flights and higher heritabilities. The flights conducted at the beginning of the 2020/2021 season (57 to 78 DAP) showed strong correlations between the indices and traits such as GY and HSW. The highest correlations were found in flights at 107, 114, and 122 DAP for TOL, NDM, and LR. In the 2019/2020 season, the flights near maturity were the most informative based on correlations, noting that no flights were conducted near flowering in that season.

### Machine Learning-Based Prediction

In this study, we employed three ML models to predict five manually measured agronomic traits in soybeans, which are associated with resistance to the stink bug complex and high yields. To make these predictions, we utilized all color, texture, and gradient histogram information extracted from UAV images. Initially, we made predictions for each flight individually, aiming to understand which flight provides the best predictions for each of the manually measured traits. In Supplementary Tables 1 and 2, we can observe the predictive capabilities of different ML models for the five manually measured traits, calculated using image-based indices.

For the regression models evaluated using the Pearson correlation metric, the AdaBoost model generally showed the best performance. The GY trait had the highest PA. In the flights conducted during the 2020/2021 season, specifically at 65, 71, and 78 DAP, predictive abilities were superior for all three models, especially with the AdaBoost model, which had correlations above 0.80. We also observed that PA decreased as flight days advanced. On the other hand, in the 2019/2020 season, the best performance was observed at 120 DAP, with a correlation of 0.7. For HSW in the 2019/2020 season, the AdaBoost model showed the best performance at 120 and 127 DAP. In 2020/2021, the AdaBoost model again stood out, following the same pattern as GY, especially at 65, 71, and 78 DAP, where correlations were high. The correlations for NDM and TOL indicate that the MLP model had inferior performance in both seasons. AdaBoost stood out with the highest correlations in flights near maturation, with correlations above 0.60 in both seasons for NDM. The same was observed for TOL, which also performed better in the last flights, and the AdaBoost and SVM models showed similar results. This same pattern was observed when using MSE to evaluate the models’ PA.

For LR, classification models were used, with accuracy and precision as evaluation metrics. In general, the three models showed similar accuracy, with AdaBoost and MLP demonstrating superior results, especially in flights close to maturity. However, the PAs for this trait were relatively low, not exceeding 0.4 in accuracy. In the 2019/2020 season, the highest accuracy was found in the flight at 127 DAP, with 0.37 using the MLP model. In the 2020/2021 season, the AdaBoost model showed the highest accuracy at 122 DAP, with 0.39. When evaluating the models using the precision metric, very similar results were found, but by this metric, the AdaBoost model was superior in both seasons.

Finally, it was found that the predictions made in the 2020/2021 season exhibited greater predictive ability compared to the 2019/2020 season for GY and HSW, possibly due to less environmental influence resulting from the high stink bug infestation in the 2019/2020 season. Additionally, in the 2020/2021 season, we conducted more flights, especially near flowering, which appear more effective for these traits. However, this pattern differed for NDM, which showed predictive capabilities exceeding 0.60 in both seasons, with the highest recorded in the 2019/2020 season. The PAs for TOL and LR were similar in both seasons and, overall, were not high, especially for LR, highlighting the difficulty of predicting these traits using ML models. Figure 8 shows the flight where the models demonstrated the highest predictive capability for each manually measured trait, calculated based on person correlation for the regression models (GY, HSW, TOL and NDM) and accuracy for the classification models (LR).

**Fig. 8:**
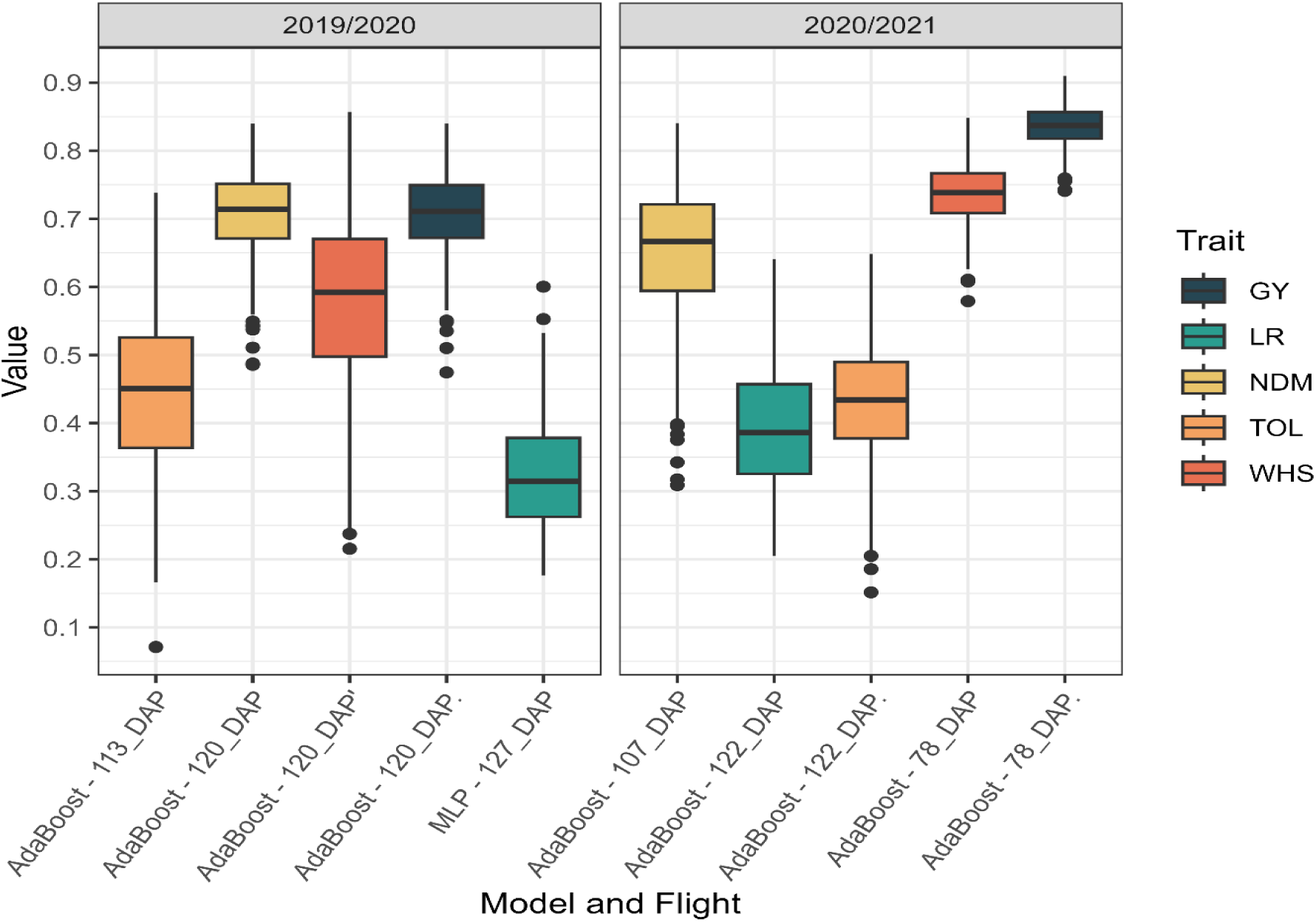
The boxplot presents the predictive ability (PA) values of the best machine learning models for each manually measured agronomic trait at different flight dates during the 2019/2020 and 2020/2021 seasons. The evaluations were performed using 10-fold cross-validation repeated 50 times, with Pearson correlation for regression models (GY, HSW, TOL, and NDM) and accuracy for classification models (LR). The flight dates and models are indicated on the x-axis, while the PA are shown on the y-axis.

We also tested different prediction scenarios by varying the combination of flights used and the prediction target year. In total, six scenarios were evaluated: two using all flights from each season (2019 and 2020), two using only the last three flights from each season, and two where data from the 2019 season were used to predict the traits in the 2020 season. These scenarios were designed to assess the predictive performance of the models based on different temporal datasets and to evaluate their ability to generalize across seasons. In Supplementary Fig. 5A and 5B, the PA of the three machine learning models (AdaBoost, MLP, and SVM) is shown for these different scenarios, with respect to the five manually measured agronomic traits (GY, LR, NDM, TOL, and HSW). This comparison highlights how different flight combinations and prediction strategies influence model performance.

In Fig. 9, we can observe that using all flights or only those close to maturity does not substantially alter the PA. However, it is evident that for most traits, there is an improvement in PA when we use all flights, instead of just one, to predict manually measured traits. For all analyzed traits, the best PA was found in the 2020/2021 season, using all flights with the AdaBoost model. Furthermore, all traits showed higher PA compared to predictions based on individual flights. The same pattern was observed in the 2019/2020 season, suggesting that using all flights can better capture temporal variations, providing a more comprehensive view of plant growth dynamics. On the other hand, as expected, using one season to predict another remains a significant challenge, even with the use of a large amount of data and machine learning models. None of the traits presented high PA values in this cross-season prediction context, reinforcing the complexity and limitations of current approaches for this type of forecasting.

**Fig. 9.**
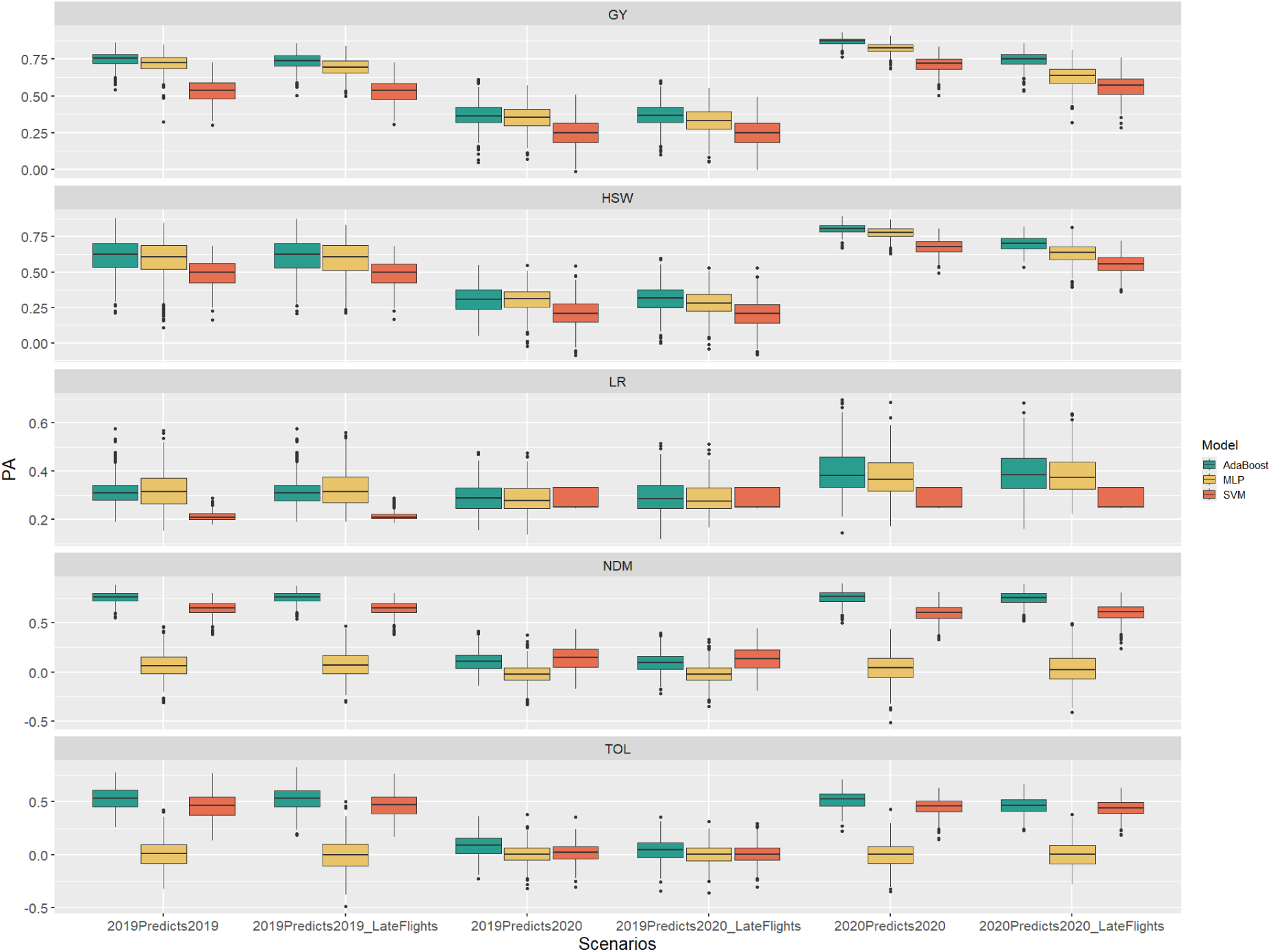
Predictive abilities (PA) of three different ML models—AdaBoost, MLP, and SVM—evaluated for the five manually measured traits across different tested scenarios. The evaluations were performed using 10-fold cross-validation repeated 50 times PAs were assessed using Pearson correlation for the regression models on traits GY, NDM, HSW, and TOL. For the classification model applied to the LR trait, accuracy was used as the evaluation metric.

Finally, we aimed to evaluate the correlations between image-based traits for the two seasons (2019/2020 and 2020/2021). In Fig. 10, the correlations of the first percentile Vari index with manually measured phenotypic traits in both seasons are presented. Interestingly, unlike the ML models, where we attempted to predict the 2020/2021 season based on the 2019/2020 data and faced challenging results, the correlations observed between seasons for some traits were quite significant. This suggests that the index has the ability to reflect consistent patterns between different seasons, at least for certain traits.

**Fig. 10:**
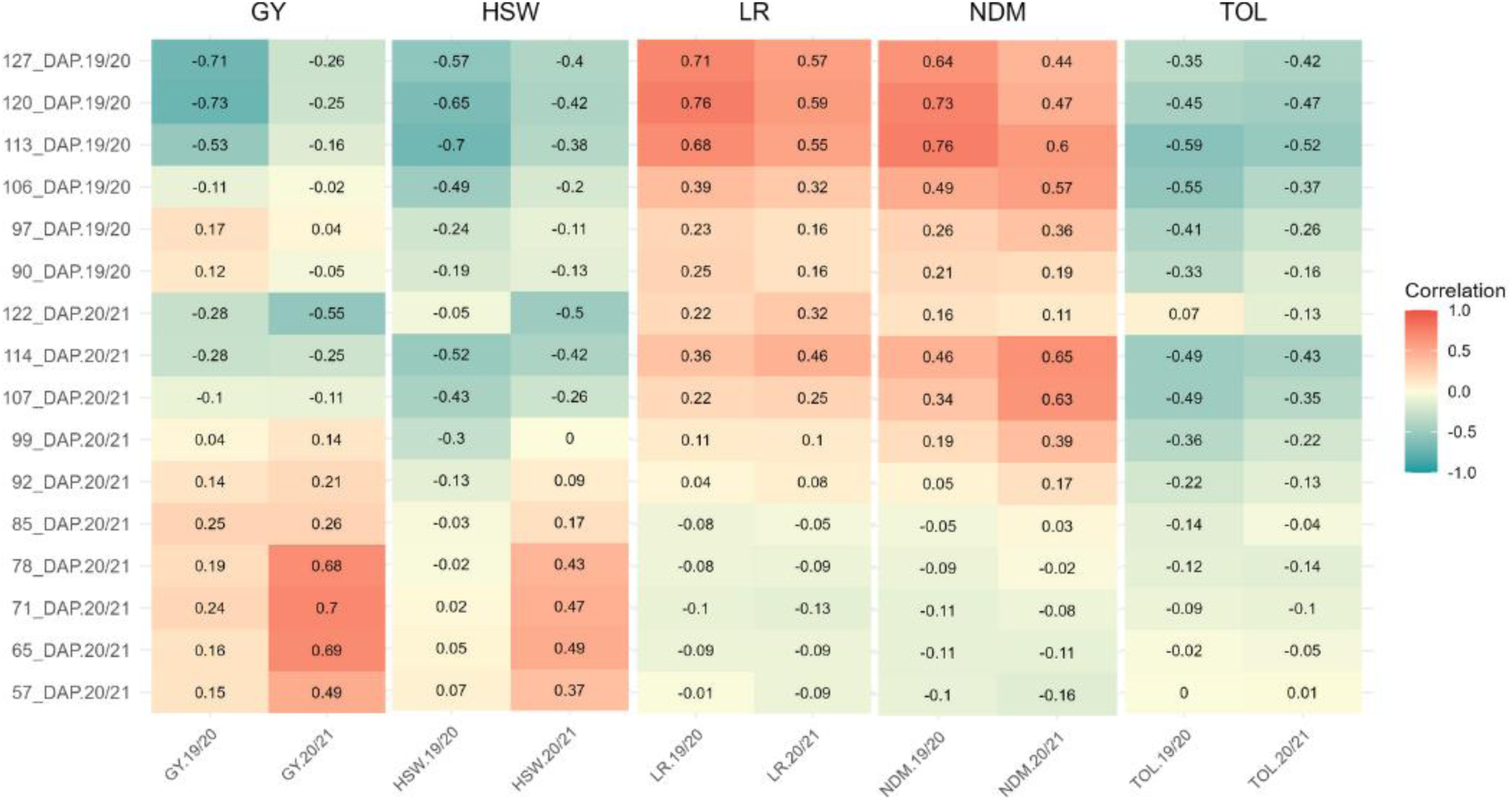
Correlations between the first percentile VARI index and the phenotypic traits GY, HSW, LR, NDM, and TOL in the 2019/2020 and 2020/2021 seasons.

Notably, the correlations between HSW and TOL in the 2019/2020 season with the VARI index in the 2020/2021 season, especially in the flights conducted at 114 and 107 days after planting (DAP), were even higher than those observed within the same season. This indicates that, for these traits, the vegetation index can capture a relevant portion of the genetic and physiological variation, despite environmental differences between the seasons. As expected, for GY, the correlations between seasons were lower, indicating that this variable is much more influenced by environmental conditions, which complicates predictions between cycles and reaffirms the complex nature of this trait.

We also conducted a decision tree analysis to understand the importance of the predictor variables. As expected, some features demonstrated significant relevance, while others exhibited low importance. Furthermore, the importance of a variable varied depending on the flight period and the year. However, certain VIs and TIs stood out, as evidenced in Supplementary Figures 6 to 10, which present the importance graph of each predictor variable in the flights that demonstrated a predictive capacity greater than 0.6 for the traits NDM, GY, and HSW. For LR e TOL, due to the low PA presented for these traits, only the most important flights were included in the graph.

For GY, the Vari_P25 stood out in the flights conducted at the end of the cycle in 2019. Additionally, other VIs also contributed significantly, along with TIs, especially ASM at 0° and 45°, and homogeneity at 45°, in the flights closer to flowering, between 57 and 78 DAP. A similar pattern was observed for the HSW characteristic, but the TIs were more prominent than the color-based VIs, particularly those at angles of 45° and 135°, in the flights near flowering, and again the Var_P25 at 122 DAP. For NDM, the Vari_P25 index was also highlighted, along with the TIs Energy at 135° and Dissimilarity at 45° in 2019, in the flights near maturation. It was also observed that some gradient histograms showed greater importance for NDM, unlike the other characteristics mentioned earlier. Finally, no variable demonstrated significant importance for LR and TOL as observed for the previous characteristics. The predictive ability of the models was influenced by all the variables collectively, with the Contrast index at 45° being the most important for LR in the last flight of 2020, while for TOL, the VARI_P25 index stands out again.

## Discussion

The use of HTP, particularly through UAVs, represents an effective tool for evaluating traits of interest in plant breeding programs. This methodology provides breeders with the ability to assess various stages of crop development and analyze a large number of genotypes across different locations, in a shorter time and with greater efficiency (Reynolds et al., 2019). In this study, we explored aerial imagery to evaluate traits of interest in soybean crops, focusing on resistance to the stink bug complex. By correlating several image descriptors with soybean traits of interest, we evaluated the feasibility of HTP in predicting traditional phenotypes with the action of stink bug attack. Additionally, we expanded such an evaluation by incorporating ML models and their capability to predict these traits using images captured by UAV. HTP protocols represent a strategy with significant potential to overcome the limitations associated with traditional large-scale phenotyping, allowing for a more detailed evaluation of traits of interest. Considering that resistance to the stink bug complex is a quantitative trait, the breeding process becomes complex, requiring extensive field evaluations (Furbank and Tester, 2011)

### Stink Bug Resistance Phenotyping in Soybean Breeding

In soybean breeding, phenotyping for resistance is well described for specific traits that condition resistance, such as pod damage, leaf retention, grain filling period, crop cycle, grain size, and the weight of healthy seeds. Rocha et al. (2014) described the latter as being strongly correlated with resistance and high yield. In our study, we observed significant correlations between traits associated with resistance, productivity, and crop cycle. These results suggest that such traits can be considered promising criteria for the simultaneous selection of high yield and resistance to stink bugs. By providing a measure of plant resistance and performance in response to stress, these traits can be used as criteria in the field selection of genotypes. We noted differences in the correlation magnitude between the two cropping seasons analyzed, likely due to varying stink bug infestations. In particular, the correlation between HSW and GY differed markedly between the seasons. The higher selection pressure in 2019/2020, contrasted with the lower pressure in 2020/2021, made selection difficult in both cases, as these situations hindered the identification of the most resistant genotypes.

We found high heritability estimates, above 0.6, for the traits analyzed, suggesting that these traits can be employed in selecting more resistant and productive genotypes. Previous studies aiming at stink bug resistance found similar results: Santos et al. (2018) identified heritabilities of 0.80 for NDM, 0.70 for GY and HSW, and 0.20 for LR in an F2:3 soybean population, while Rocha et al. (2015) reported heritabilities of 0.75 for NDM, 0.24 for GY, 0.35 for HSW, and 0.67 for LR in a RIL population, both originating from the cross between IAC 100 (resistant) and CD 215 (susceptible) under natural stink bug infestation. The differences in heritabilities observed between studies can be attributed to several factors. GY and HSW are quantitative traits that are highly influenced by varying environmental conditions, leading to different heritability estimates under different conditions (Han et al., 2012). For LR, its value heavily depends on the evaluator’s assessment, making it challenging to consistently achieve high heritability estimates (Godoi and Pinheiro, 2009; Pinheiro et al., 2005). Additionally, the genetic background of the populations studied, and the specific environmental conditions (Aditya et al., 2011; Li et al., 2020).

Among the traits studied, we observed that genetic variance plays a significant role in the total variation found, indicating the presence of variability in the evaluated population and enabling the selection of genotypes. This variability is influenced by the parental lines involved in the crosses, which are quite diverse, including IAC 100, known for its multiple mechanisms of stink bug resistance (McPherson et al., 2007; Veiga et al., 1999). The first step for the success of a breeding program is the careful selection of parents, aiming for genetic diversity and the identification of combinations capable of generating superior genotypes while preserving variability (Falconer et al., 1996). However, for many years, the focus of breeding has primarily been on high yields, making crosses that do not always contribute to an increase in variability. This can hinder the long-term success of the program, as it limits genetic progress for stink bug resistance (Bernardo, 2010).

Despite these traits being promising for use in breeding, visual assessment becomes impractical and inaccurate on a large scale, a common challenge encountered in plant breeding programs, especially when evaluating insect resistance (Tang and He, 2021). Although the use of images in field conditions for soybean has been explored in various applications, the use of this tool for evaluating insect resistance, particularly stink bugs, is still scarcely documented in the literature (Iost-Filho et al., 2022; Marston et al., 2020, 2022). Through imaging, it is possible to calculate many features, such as VIs, which have a biological basis and allow for the quantitative assessment of physiological processes using reflected wavelengths. Additionally, TIs capture structural and spatial information about plant canopies, reflecting aspects like leaf arrangement and density (Araus and Cairns, 2014; Li et al., 2014).

### Image-based Phenotyping for Stink Bug Resistance

Given the utility of aerial images, their application extends to evaluating a wide range of traits in plant breeding programs. VIs are measures related to photosynthetic activity and the physiological state of plants, especially concerning their response to stress and overall health (Krause et al., 2020). These indices can capture subtle differences in plant composition and photosynthetic activity, which may be influenced by different environmental conditions and biotic stresses (Kefauver et al., 2015). Additionally, VIs are used in many studies for predicting agricultural productivity due to their stable and superior performance (Ballester et al., 2017; Zhou et al., 2017). Therefore, variations in VIs among different tested genotypes can provide insights into how plants respond and adapt to different stresses. In our study, we identified promising VIs that can be used individually in predicting traits of interest. These VIs not only exhibited high estimates of heritability and accuracy at different flight times but also revealed highly significant correlations, particularly the correlations found between traits associated with resistance in flights near maturity and harvest, especially in environments stressed by high insect incidence.

Using VIs derived from RGB images captured by UAV and visual measurements to quantify senescence in wheat, Li et al. (2023) identified that genotypes with slow senescence were effectively selected. In a study conducted by Resende et al. (2024) in maize, flights conducted at the V5 and V8 phenological stages showed the highest correlations between the VI GLI and productivity. However, they also observed that these correlations can be highly variable depending on environmental conditions and plant phenological stages. Additionally, it is worth noting that we used the first percentile of the pixels instead of the average, as is done in many other studies, because it showed a higher correlation with the traits of interest. This is likely because stink bug infestation is uneven in the field and, consequently, within the plot (Fernandes et al., 2019). Thus, considering only the first quartile of the pixels ensures that we are evaluating those plants that were attacked by stink bugs. This method allows for a more accurate assessment of resistance to stink bug attacks and can be an interesting strategy for breeding programs focused on insect resistance.

In soybeans, Bai et al. (2022) used RGB images to make yield predictions, especially under lodging conditions, and found better predictive abilities at 48 days after planting, near the flowering stage. This underscores the importance of conducting flights at different plant developmental stages to capture critical phenological changes that influence yield potential. Additionally, these authors highlighted that texture information from the images has a high potential to contribute to yield estimation. Using different sensors coupled with UAVs, Maimaitijiang et al. (2020) reported that the inclusion of texture information from soybean canopies improved the accuracy of yield prediction, with R² ranging from 0.65 to 0.72. TIs represent spatial variations in color intensity in images and can capture repetitive patterns that are useful for assessing aspects such as canopy uniformity, which are associated with plant vigor and, consequently, productivity (Ma et al., 2022).

In our study, we observed that VIs, particularly VARI_P25, not only presented higher heritabilities but also showed higher and more consistent correlations with the traits of interest. This suggests that, while TIs can be useful, VIs provide a more direct and effective assessment of the physiological conditions of the plants and their response to stress, especially under stink bug infestation conditions. Additionally, the PCA based on TIs, despite showing lower correlations and heritabilities, also proved to be interesting, particularly for the trait TOL in the 2019/2020 season, with a correlation of 0.58, higher than the other VIs. This indicates that while VIs are generally more effective, TIs can provide valuable information in specific contexts, complementing the analysis of stink bug resistance and other agronomic traits.

Our results showed that the selected indices are strongly associated with visual characteristics, suggesting that these indices can be used individually for selection. However, understanding how these indices can contribute to the prediction of a particular characteristic when used together is crucial. In this context, the use of ML models can be adopted to explore and maximize all the information contained in complex datasets, such as those generated by images. Therefore, we evaluated different ML-based models, employing the 124 features obtained related to color, texture, and vegetation indices. In this analysis, we tested not only individual flight times but also different prediction scenarios to understand how the combined flight times can contribute to greater accuracy.

### Machine Learning Prediction of Stink Bug Resistance

Many soybean studies have utilized images obtained by UAVs and different ML models to address common challenges in crop improvement. Hassanijalilian et al. (2020) tested different decision tree-based models to classify iron deficiency chlorosis in 40 soybean cultivars using RGB images. The authors found that the AdaBoost model exhibited the best performance contrasted to random forest (RF), and decision tree, with an average f1-score of 0.75. Additionally, RF models, also based on decision trees, showed high accuracy in distinguishing soybean genotypes resistant and susceptible to dicamba, with overall accuracies ranging from 0.69 to 0.75, like this artificial neural network (Vieira et al., 2022). It is important to note that LR, like the symptoms of iron deficiency chlorosis and dicamba phytotoxicity, visibly manifests through chlorophyll loss in the leaves. In our study, we observed that the AdaBoost model also produced promising results in predicting manually measured traits, both when using individual flights and when using combined flight times, suggesting that decision tree-based models can be effective in predicting these characteristics. However, we found PAs much lower than those reported by these authors. One hypothesis is that in the 2019/2020 season, the attack was extremely severe, resulting in many scores of 5, while in the following season, there were almost no symptoms in the field, resulting in few scores ranging from 1 to 5. Thus, the model had difficulty adequately distinguishing the symptoms due to the lack of variation in the data within the same season. These observations suggest that an ideal control level for stink bugs should be established to ensure a more balanced and accurate assessment of resistance traits.

For GY and HSW, we found that the best times for prediction are those closest to flowering, although good prediction capabilities are also found in other flights. Zhou et al. (2020) found similar results, using a convolutional neural network (CNN) model to estimate soybean productivity under water stress with RGB and multispectral image features, as well as visual characteristics. Their model explained 78% of the feature, indicating that images collected in the early and late stages of reproductive growth are promising for productivity estimation.

By incorporating all flights to predict GY, the results indicate that a greater number of flights contributes to capturing a higher degree of phenotypic and environmental variability. Flights conducted close to the flowering period can capture significant environmental variations, such as incomplete plant development or establishment failures, which may interfere with data interpretation without necessarily reflecting the true genetic nature of the trait being evaluated. Additionally, stink bug infestations, which directly affect soybeans during the R5 reproductive stage, considered critical for infestation, can be more adequately monitored by flights conducted near the maturation period when pest damage becomes visible. These flights, therefore, more accurately capture genetic information, providing a more precise estimate of genetic performance, unlike flights conducted during the flowering stage, which may not fully capture the genetic effects associated with yield.

Although UAVs are widely used in plant breeding, their use in the field for stink bug phenotyping is not reported. However, some studies have explored this tool to identify aphid infestation symptoms in soybeans. Using SVM models for classification, the stress caused by soybean aphids was effectively identified using hyperspectral reflectance, achieving up to 89.4% accuracy with optimal wavelength combinations (Marston et al., 2022). Furthermore, using data obtained from multispectral cameras and simple linear regression analyses, it was possible to identify soybean aphid-induced stress (Marston et al., 2020). However, these studies were conducted with controlled infestation in cages and using few or only one genotype, conditions that differ from those found in breeding programs. Despite these limitations, the results demonstrate the potential of this tool for application under field conditions and the possibility of using other cameras, such as hyperspectral and multispectral, which can provide new information and improve prediction.

The results of our study revealed that, in general, the best PAs were achieved in the 2020/2021 season for the traits HSW and GY. However, caution should be exercised when selecting individuals based on HSW during periods of low stink bug infestation, as this may lead to the selection of more productive individuals but not necessarily more resistant ones. This is because this trait showed a highly significant correlation with GY in this season. For NDM, which also showed high PA, the best results were obtained in the 2019/2020 season (0.71). However, TOL and LR showed lower PAs in both seasons. The high stink bug infestation in the 2019/2020 season resulted in greater selection pressure, making it difficult to identify more resistant genotypes and compromising the accuracy of prediction models, especially for resistance-related traits. Establishing an optimal level of stink bug control is a challenge faced by breeding programs, and this discussion is crucial for developing more precise and effective selection strategies. As observed for GY and HSW, the use of a greater number of flights for the other traits also allows for an increase in PA, even though this increment is modest. Expanding data collection through additional flights improves the accuracy of prediction models by providing a more comprehensive estimate of genotype-environment interactions, resulting in a more accurate and efficient selection of genotypes with superior genetic performance.

Crop prediction in plant breeding programs using data from previous years can be a valuable tool, but it presents considerable challenges, mainly due to environmental variations between seasons (Crossa et al., 2017; Yang et al., 2017). This is particularly evident when UAV-acquired images are used for this task, as the different climatic and management conditions encountered in each flight affect the accuracy of the predictions. This challenge was observed in the difficulty of obtaining high PAs when attempting to predict agronomic traits of one season using data from the previous season, underscoring the complexity of establishing this strategy in breeding programs (Araus et al., 2018; Gracia-Romero et al., 2019).

However, when analyzing the correlations between the first percentile VARI vegetation index and manually measured phenotypic traits, we identified significant correlations, although not extremely high. These correlations, though modest, can be useful for an initial genotype screening, allowing the exclusion of less promising materials and consequently optimizing resources and evaluation time in subsequent seasons (Zhou et al., 2022). In particular, the correlations observed between HSW and TOL in the 2019/2020 season with the VARI index from the 2020/2021 season were higher than those observed within the same season. This suggests that, for these traits, the vegetation index captures a relevant portion of genetic and physiological variation, even in the face of environmental differences between crop cycles. On the other hand, the correlations for GY were significantly lower between seasons, indicating that this variable is much more influenced by environmental conditions, corroborating its complex and quantitative nature.

We did not identify consistent patterns among the most important indices in the decision tree analysis. Each flight and each characteristic exhibited different patterns of influence from the indices used. However, one highlight was the VIs Vari_25Percent. This index was present among the flights with the highest predictive capacity, except for LR, where it was not as influential. Nevertheless, for the other traits, Vari_25Percent proved to be crucial for predictive capacity. Additionally, the VARI_25Percent index obtained high correlations when evaluated individually in relation to manually measured traits. Vari is an index based on the variability of plant vigor and stress, measured by its greenness (Gitelson et al., 2002). It is used to interpret various vegetation factors, such as grain weight under stress conditions (Rodene et al., 2022), yield (Silva et al., 2022), and leaf area (Hasan et al., 2019; Zhang et al., 2019). TIs also proved to be important in ML models for all traits, with emphasis on the indices calculated at angles of 45° and 135°.

This work is pioneering in utilizing UAVs and ML for evaluating stink bug resistance in breeding programs, offering insights that fuel discussions on the use of these tools in soybean phenotyping. However, it also underscores the need for further studies to optimize stink bug resistance selection. Continued research on this issue, including the evaluation of other methodologies and the incorporation of new technologies, promises significant advances in soybean genetic breeding for stink bug resistance.

## Statements and Declarations

### CRediT Author Contribution Statement

MO: Conceptualization, Methodology, Software, Validation, Formal analysis, Investigation, Data Curation, Writing - Original Draft, Writing - Review & Editing, Visualization. AA: Methodology, Software, Validation, Formal analysis, Investigation, Data Curation, Writing - Review & Editing, Visualization. PB: Conceptualization, Methodology, Investigation, Data Curation. AM: Conceptualization, Methodology, Investigation, Data Curation. FC: Data Curation. FK: Data Curation. FF: Data Curation. JPazini: Data Curation, Writing - Review & Editing. PY: Validation, Data Curation, Resources, Supervision. JPinheiro: Conceptualization, Validation, Investigation, Data Curation, Writing - Review & Editing, Resources, Supervision, Project administration, Funding acquisition

### Data Availability

Data will be made available on request.

## Acknowledgments

This work was supported by grants from the Conselho Nacional de Desenvolvimento Científico e Tecnológico (CNPq), Fundação de Amparo à Pesquisa do Estado de São Paulo (FAPESP), and the Coordenação de Aperfeiçoamento de Pessoal de Nível Superior (CAPES – Financial Code 001). AA received a PhD fellowship from FAPESP (2019/03232-6).

## SUPPLEMENTARY RESULTS

**Supplementary Table 1.** Predictive abilities (PA) of three different ML models, AdaBoost, MLP, and SVM, for the five manually measured traits in two growing seasons, 2019/2020 and 2020/2021, at each flight time. PAs were evaluated using Pearson correlation for regression models on the traits GY, NDM, HSW, and TOL. For classification models, used for the LR trait, accuracy was the metric used.

|  |  | GY |  |  | HSW |  |  | NDM |  |  | TOL |  |  | LR |  |  |
| --- | --- | --- | --- | --- | --- | --- | --- | --- | --- | --- | --- | --- | --- | --- | --- | --- |
| Year | DAP | AdB. | MLP | SVM | AdB. | MLP | SVM | AdB. | MLP | SVM | AdB. | MLP | SVM | AdB. | MLP | SVM |
| 2019/<br>2020 | 90 | 0.46 | 0.12 | 0.21 | 0.18 | 0.12 | 0.1 | 0.29 | 0 | 0.2 | 0.11 | 0.01 | 0.11 | 0.22 | 0.22 | 0.20 |
|  | 97 | 0.44 | 0.25 | 0.26 | 0.12 | 0.15 | 0.12 | 0.3 | -0.1 | 0.23 | 0.14 | -0.03 | 0.06 | 0.20 | 0.23 | 0.20 |
|  | 106 | 0.51 | 0.4 | 0.32 | 0.43 | 0.32 | 0.25 | 0.6 | 0.09 | 0.38 | 0.41 | 0 | 0.3 | 0.29 | 0.23 | 0.20 |
|  | 113 | 0.6 | 0.42 | 0.37 | 0.51 | 0.47 | 0.39 | 0.67 | 0.03 | 0.53 | 0.47 | 0.08 | 0.39 | 0.27 | 0.22 | 0.20 |
|  | 120 | 0.7 | 0.58 | 0.56 | 0.59 | 0.56 | 0.47 | 0.71 | 0.05 | 0.6 | 0.44 | 0.12 | 0.44 | 0.34 | 0.33 | 0.21 |
|  | 127 | 0.64 | 0.56 | 0.55 | 0.55 | 0.54 | 0.49 | 0.62 | 0.05 | 0.54 | 0.45 | -0.06 | 0.4 | 0.36 | 0.37 | 0.22 |
| 2020/<br>2021 | 57 | 0.76 | 0.64 | 0.68 | 0.69 | 0.57 | 0.6 | 0.37 | 0.06 | 0.27 | 0.23 | 0 | 0.21 | 0.29 | 0.29 | 0.28 |
|  | 65 | 0.81 | 0.64 | 0.69 | 0.72 | 0.59 | 0.61 | 0.34 | 0 | 0.26 | 0.21 | -0.03 | 0.24 | 0.29 | 0.29 | 0.28 |
|  | 71 | 0.82 | 0.64 | 0.68 | 0.71 | 0.57 | 0.58 | 0.31 | 0.01 | 0.21 | 0.17 | 0.05 | 0.16 | 0.31 | 0.31 | 0.28 |
|  | 78 | 0.84 | 0.63 | 0.67 | 0.73 | 0.59 | 0.57 | 0.24 | -0.1 | 0.13 | 0.25 | -0.01 | 0.16 | 0.31 | 0.30 | 0.28 |
|  | 85 | 0.78 | 0.44 | 0.34 | 0.69 | 0.42 | 0.3 | 0.31 | -0.1 | 0.26 | 0.14 | 0 | 0.1 | 0.32 | 0.29 | 0.28 |
|  | 92 | 0.74 | 0.43 | 0.34 | 0.66 | 0.43 | 0.34 | 0.44 | -0.2 | 0.35 | 0.24 | 0.01 | 0.23 | 0.30 | 0.29 | 0.28 |
|  | 99 | 0.72 | 0.24 | 0.21 | 0.61 | 0.28 | 0.24 | 0.51 | -0.1 | 0.42 | 0.24 | 0.04 | 0.25 | 0.29 | 0.29 | 0.28 |
|  | 107 | 0.68 | 0.41 | 0.33 | 0.63 | 0.43 | 0.37 | 0.64 | 0.12 | 0.54 | 0.36 | -0.03 | 0.3 | 0.32 | 0.30 | 0.28 |
|  | 114 | 0.63 | 0.42 | 0.42 | 0.58 | 0.47 | 0.47 | 0.56 | -0.2 | 0.51 | 0.36 | -0.03 | 0.32 | 0.35 | 0.33 | 0.28 |
|  | 122 | 0.65 | 0.51 | 0.49 | 0.61 | 0.5 | 0.43 | 0.61 | 0.18 | 0.48 | 0.44 | 0.01 | 0.43 | 0.39 | 0.36 | 0.28 |

**Supplementary Table 2.** Predictive abilities (PA) of three different ML models, AdaBoost, MLP, and SVM, for the five manually measured traits in two growing seasons, 2019/2020 and 2020/2021, at each flight time. PAs were evaluated using mean squared error for regression models on the traits GY, NDM, HSW, and TOL. For classification models, used for the LR trait, precision was the metric used.

|  |  | GY |  |  | HSW |  |  | NDM |  |  | TOL |  |  | LR |  |  |
| --- | --- | --- | --- | --- | --- | --- | --- | --- | --- | --- | --- | --- | --- | --- | --- | --- |
| Year | DA<br>P | AdB. | MLP | SVM | AdB. | MLP | SVM | AdB | MLP | SVM | Ad<br>B | MLP | SVM | AdB. | MLP | SVM |
| 2019.<br>2020 | 90 | 270181 | 307498 | 360460 | 28776 | 24578 | 29388 | 10.1 | 80.4 | 13.2 | 0.00<br>54 | 4.8 | 0.006<br>1 | 0.21 | 0.19 | 0.08 |
|  | 97 | 236327 | 305725 | 358940 | 27563 | 22077 | 28270 | 8.3 | 122.3 | 11.5 | 0.00<br>54 | 48.1 | 0.005<br>9 | 0.19 | 0.23 | 0.09 |
|  | 106 | 189110 | 243736 | 355585 | 22829 | 18643 | 28192 | 7.4 | 204.2 | 10.5 | 0.00<br>57 | 54.2 | 0.005<br>9 | 0.26 | 0.24 | 0.09 |
|  | 113 | 217567 | 250363 | 360064 | 24405 | 19658 | 28818 | 9.6 | 98.5 | 11.4 | 0.00<br>54 | 8.5 | 0.006<br>0 | 0.27 | 0.23 | 0.09 |
|  | 120 | 290920 | 442840 | 364307 | 40760 | 26454 | 29818 | 14.6 | 138.5 | 14.4 | 0.00<br>71 | 3.9 | 0.006<br>5 | 0.30 | 0.29 | 0.16 |
|  | 127 | 296165 | 358319 | 363776 | 34329 | 26219 | 29701 | 14.6 | 85.5 | 14.2 | 0.00<br>68 | 7.3 | 0.006<br>5 | 0.34 | 0.32 | 0.18 |
| 2020.<br>2021 | 57 | 475925 | 722001 | 867394 | 386777 | 498260 | 611058 | 7.6 | 86.3 | 11.7 | 0.01<br>49 | 2.5 | 0.015<br>3 | 0.28 | 0.29 | 0.18 |
|  | 65 | 530779 | 727973 | 865350 | 406265 | 483693 | 608497 | 9.5 | 2184 | 11.0 | 0.01<br>48 | 3518 | 0.015<br>2 | 0.28 | 0.28 | 0.18 |
|  | 71 | 508380 | 654901 | 855257 | 391632 | 467282 | 607178 | 8.6 | 285 | 12.1 | 0.01<br>37 | 25.0 | 0.013<br>8 | 0.30 | 0.24 | 0.18 |
|  | 78 | 364475 | 529435 | 858763 | 330987 | 424552 | 608339 | 12.0 | 1029 | 12.8 | 0.01<br>62 | 74.0 | 0.016<br>2 | 0.30 | 0.24 | 0.18 |
|  | 85 | 296048 | 543537 | 848846 | 298452 | 415293 | 600972 | 12.2 | 259 | 12.9 | 0.01<br>64 | 24.5 | 0.016<br>0 | 0.34 | 0.28 | 0.18 |
|  | 92 | 279382 | 563397 | 860841 | 314961 | 439028 | 609376 | 12.4 | 274 | 13.1 | 0.01<br>66 | 61.1 | 0.016<br>6 | 0.30 | 0.29 | 0.18 |
|  | 99 | 261983 | 564648 | 862498 | 294017 | 419245 | 610626 | 13.6 | 332 | 13.2 | 0.01<br>60 | 21.1 | 0.016<br>4 | 0.29 | 0.28 | 0.18 |
|  | 107 | 351000 | 702333 | 868343 | 324166 | 507863 | 614002 | 12.4 | 173 | 13.1 | 0.01<br>67 | 2.9 | 0.016<br>7 | 0.32 | 0.25 | 0.18 |
|  | 114 | 40551 | 713408 | 866994 | 359308 | 502075 | 612688 | 10.8 | 144 | 13.0 | 0.01<br>59 | 2.0 | 0.016<br>0 | 0.34 | 0.31 | 0.18 |
|  | 122 | 433084 | 885817 | 870498 | 399652 | 597425 | 614897 | 9.8 | 1099 | 12.8 | 0.01<br>61 | 68.1 | 0.015<br>8 | 0.40 | 0.33 | 0.18 |

**Supplementary Fig. 1.**
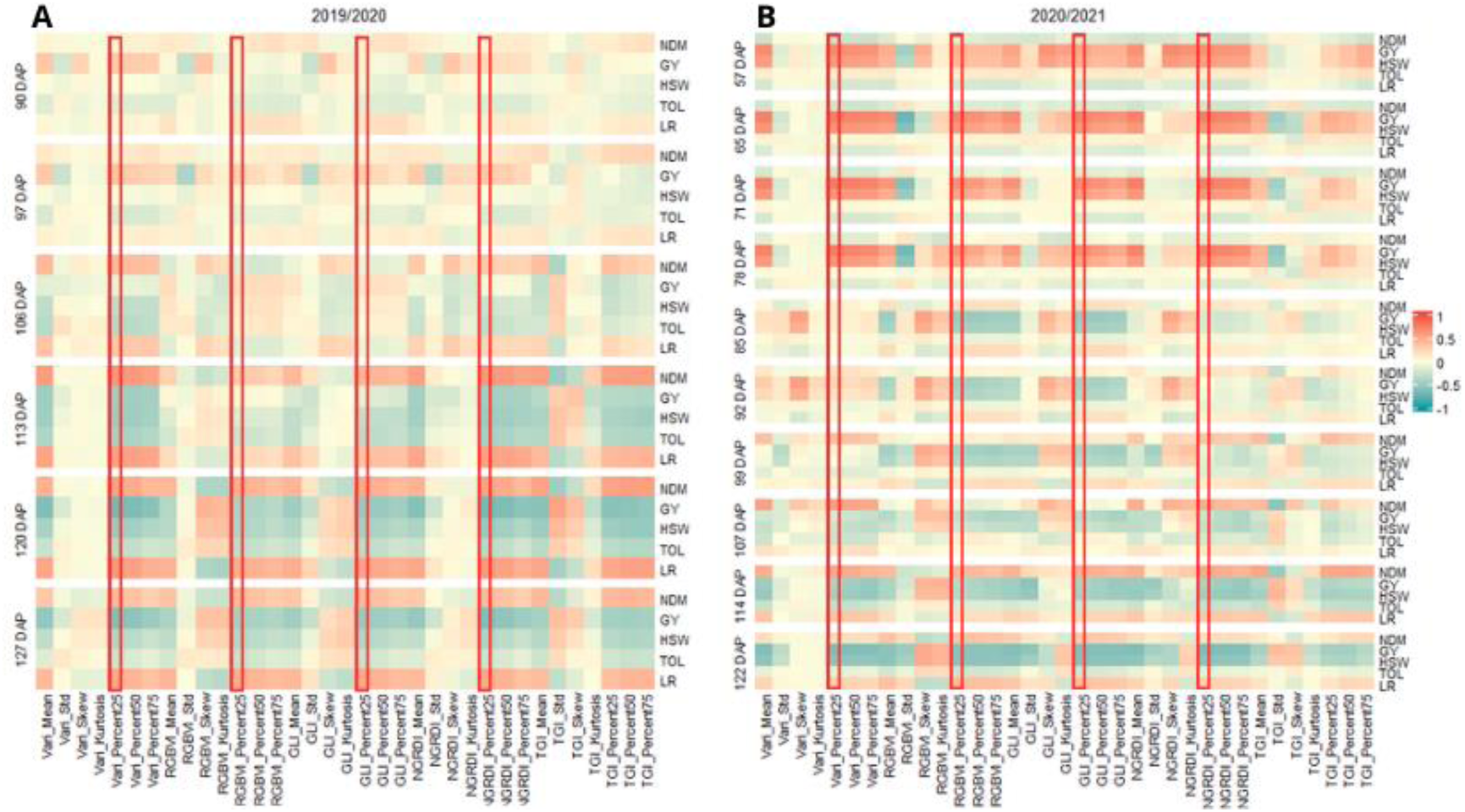
Heatmap displaying the correlation between manually measured traits and vegetation indices across different phenological stages for two consecutive growing seasons, 2019/2020 (A) and 2020/2021 (B). Traits assessed include NDM (Number of Days to Maturity), GY (Grain Yield), HSW (Healthy Seed Weight), TOL (Tolerance), and LR (Leaf Retention). The red rectangles highlight vegetation indices selected based on the 25th percentile (Percent25), marking the most relevant indices for further analysis.

**Supplementary Fig. 2.**
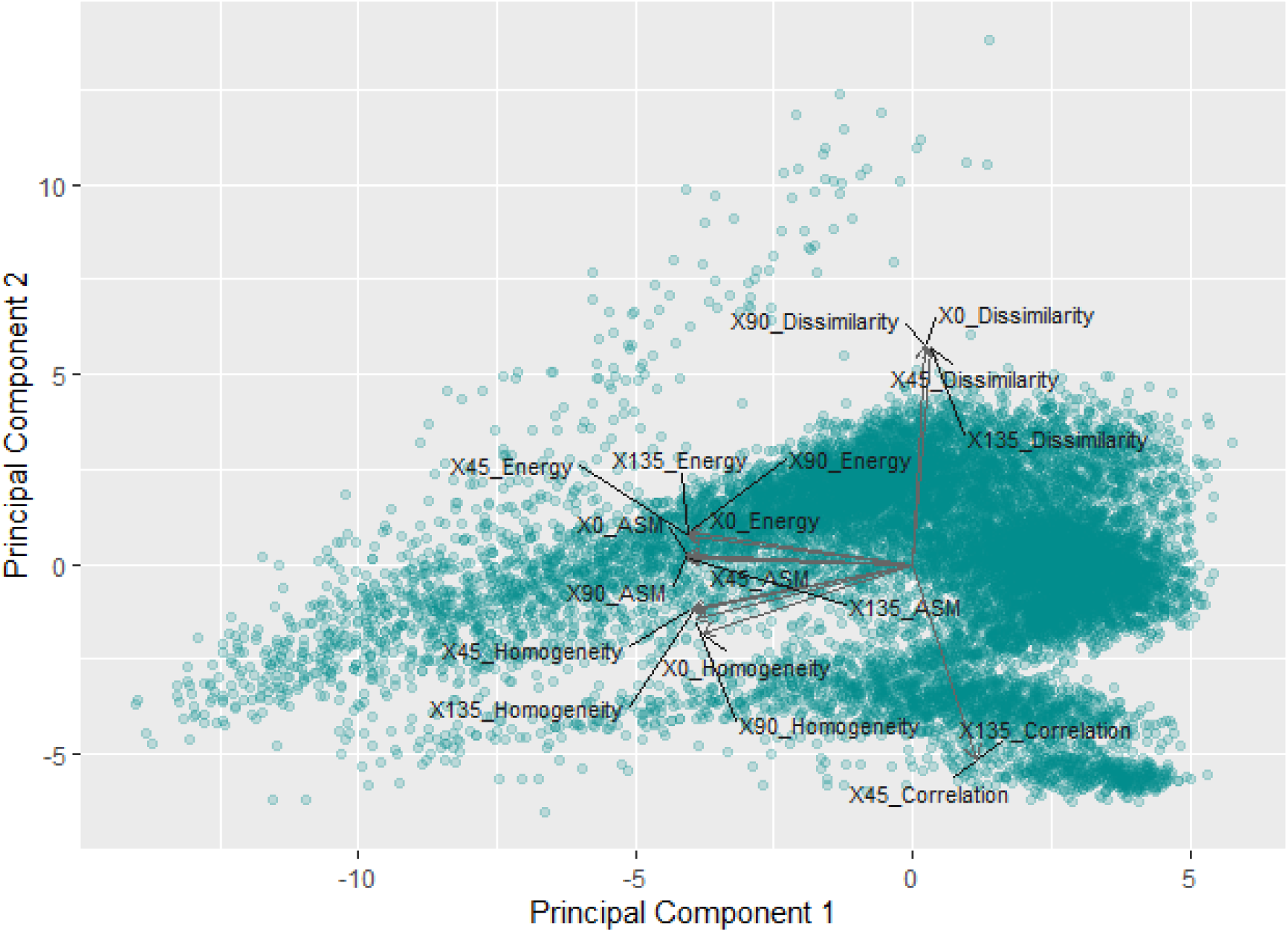
Principal Component Analysis (PCA) biplot illustrating the distribution of texture indices based on the first two principal components. The vectors represent different texture indices, including Dissimilarity, Energy, Homogeneity, ASM (Angular Second Moment), and Correlation, calculated at four angles (0°, 45°, 90°, and 135°). The scatter points represent individual observations, with each point corresponding to a plot in the field, recorded at different flight times and across multiple growing seasons.

**Supplementary Fig. 3.**
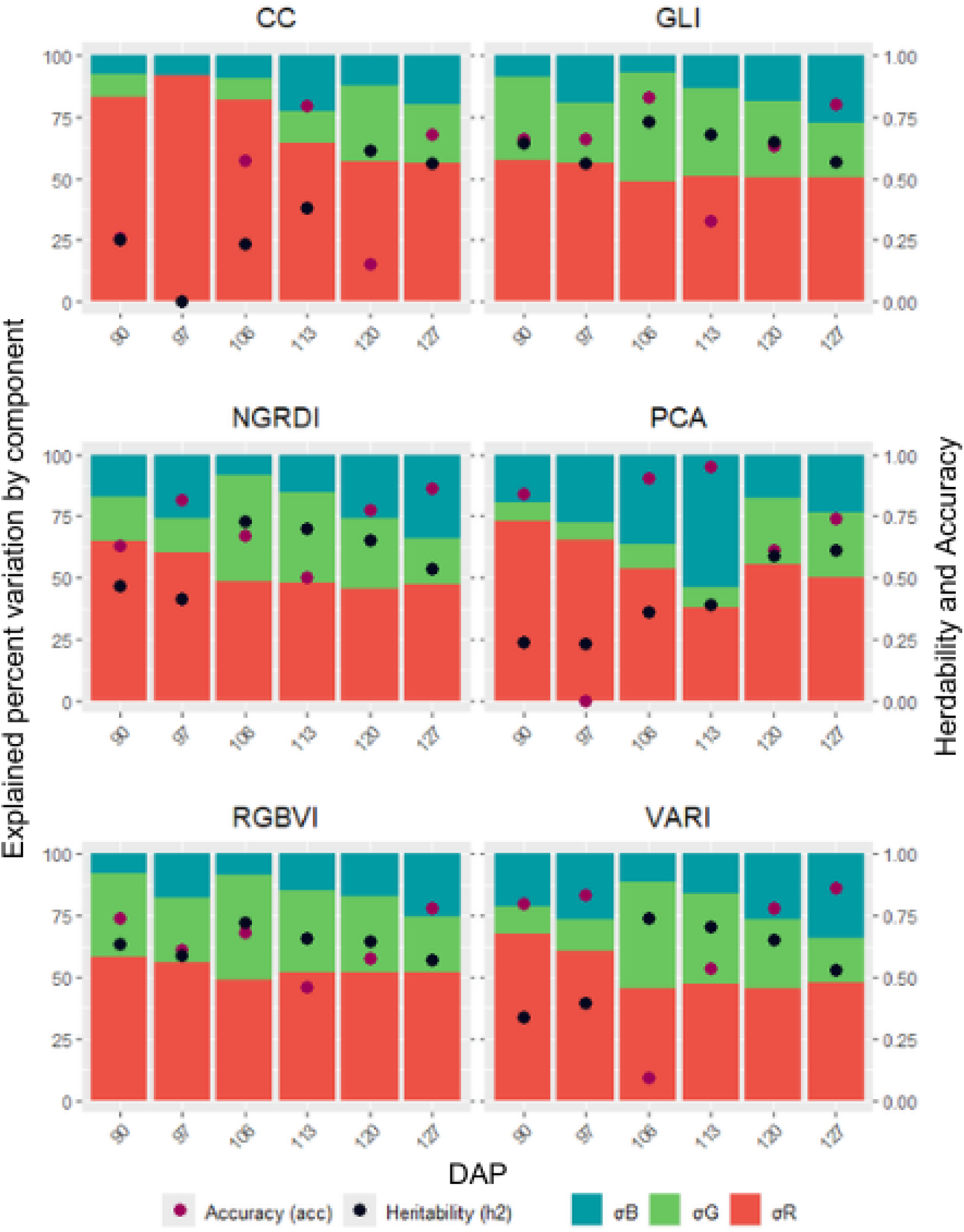
Percentage of variation explained by variance components, heritability, and accuracy for image-derived traits during the 2019/2020 growing season for each flight.

**Supplementary Fig. 4.**
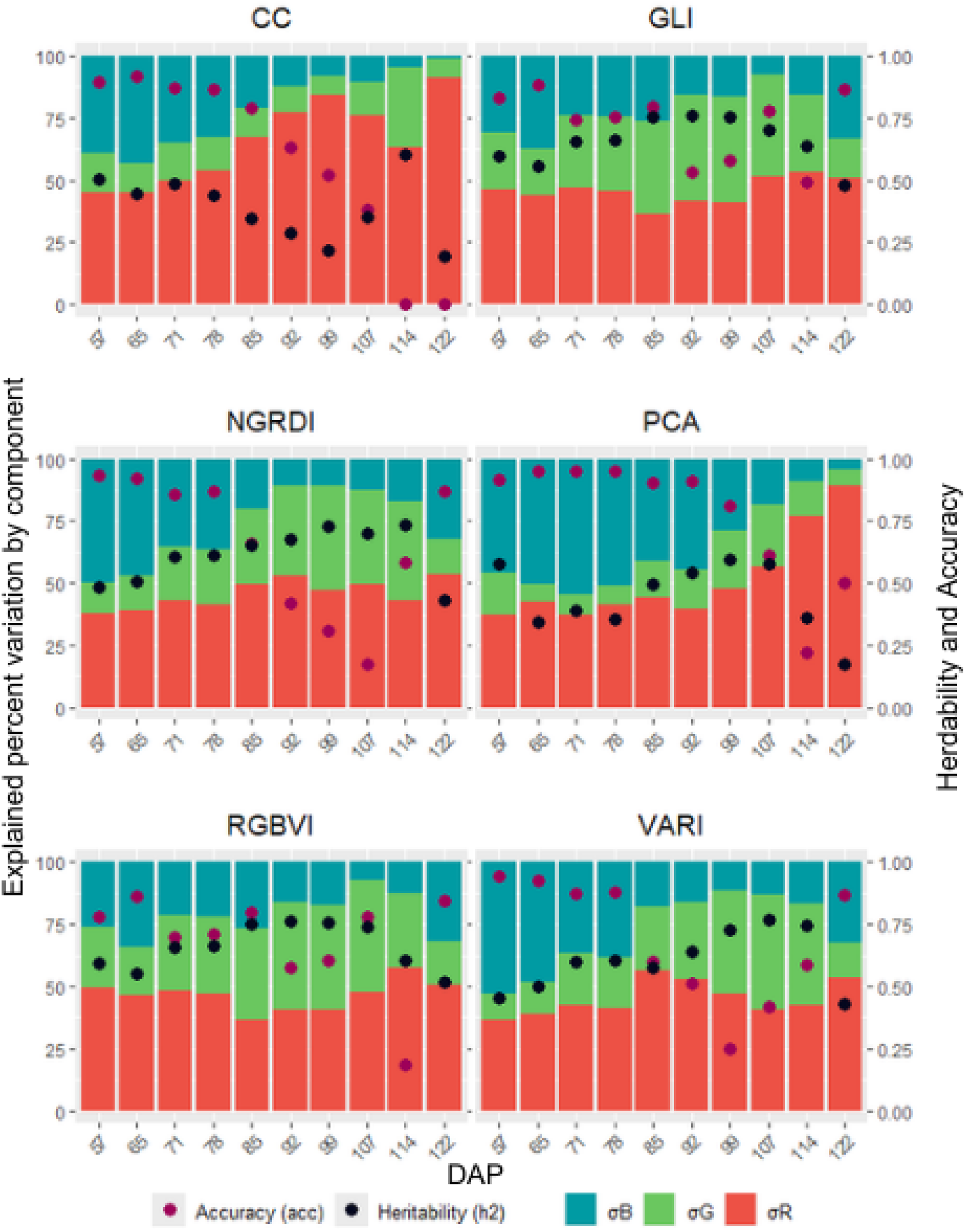
Percentage of variation explained by variance components, heritability, and accuracy for image-derived traits during the 2020/2021 growing season for each flight.

**Supplementary Fig. 5.**
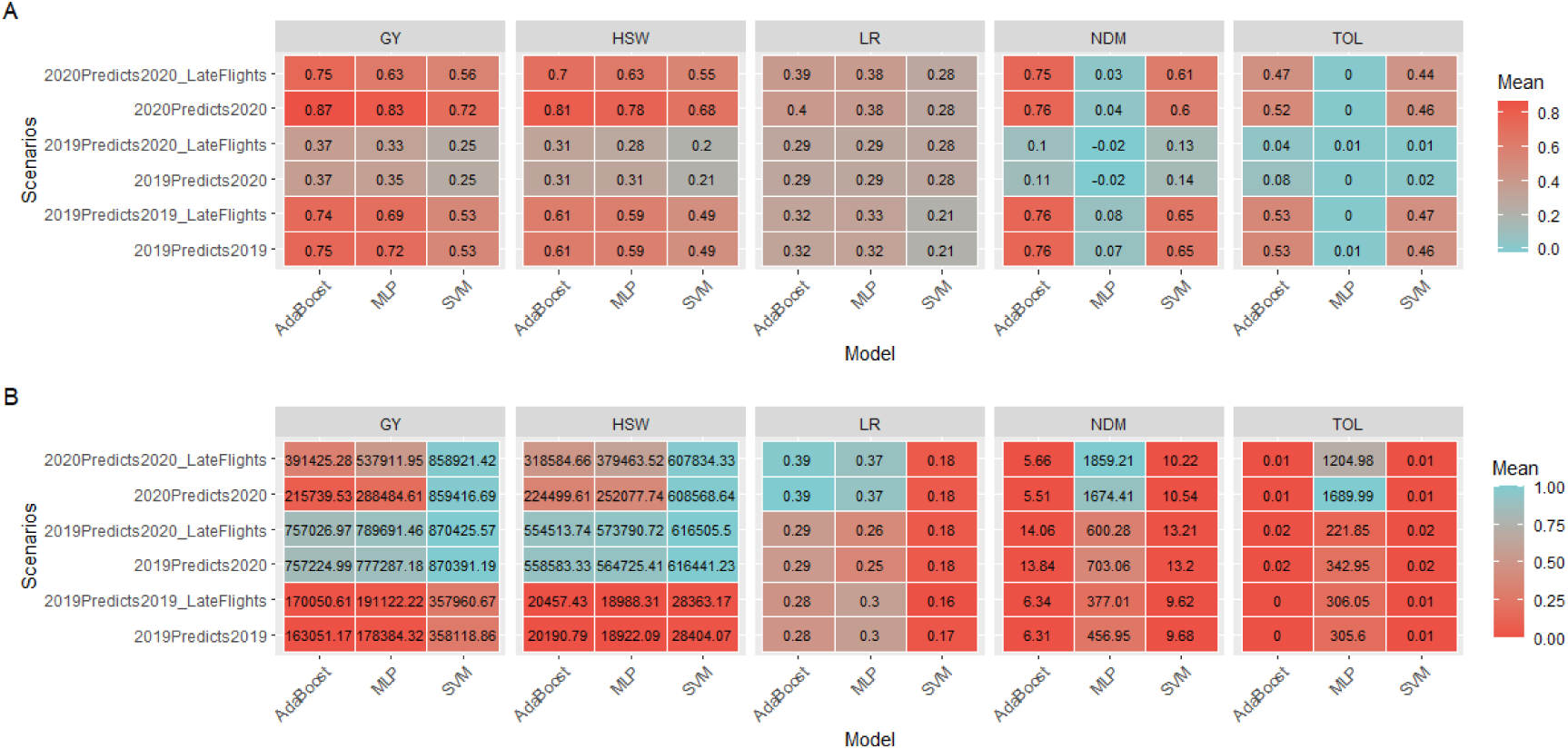
Predictive abilities (PA) of three different ML models—AdaBoost, MLP, and SVM— evaluated for the five manually measured traits across different tested scenarios. The evaluations were performed using 10-fold cross-validation repeated 50 times. **A.** PAs were assessed using Pearson correlation for the regression models on traits GY (Grain Yield), NDM (Number of Days to Maturity), HSW (Healthy Seed Weight), and TOL (Tolerance). For the classification model applied to the LR (Leaf Retention) trait, accuracy was used as the evaluation metric. **B.** PAs were assessed using mean squared error (MSE) for the regression models on traits GY, NDM, WHS, and TOL. For the classification model applied to LR, precision was used as the evaluation metric.

**Supplementary Fig. 6.**
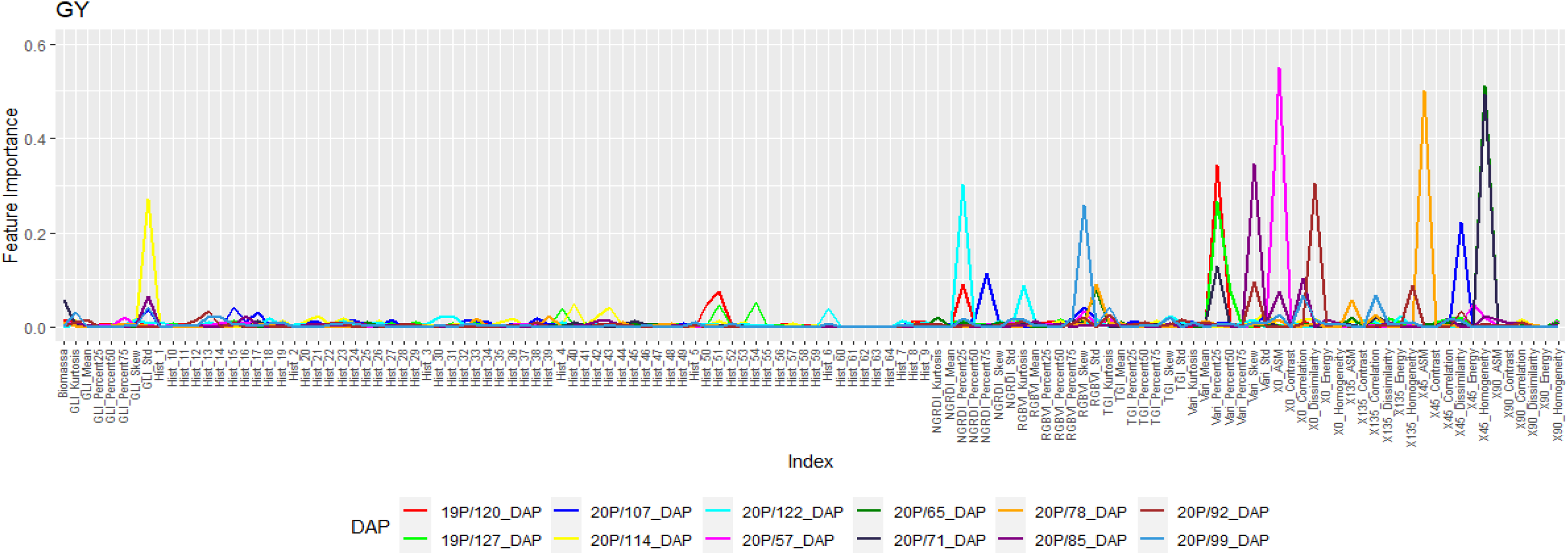
Importance of each predictor variable for predicting GY in each flight where the predictive ability was greater than 0.60. The importance was calculated based on decision tree analysis, estimating the importance of each image-derived feature using the Gini index.

**Supplementary Fig. 7.**
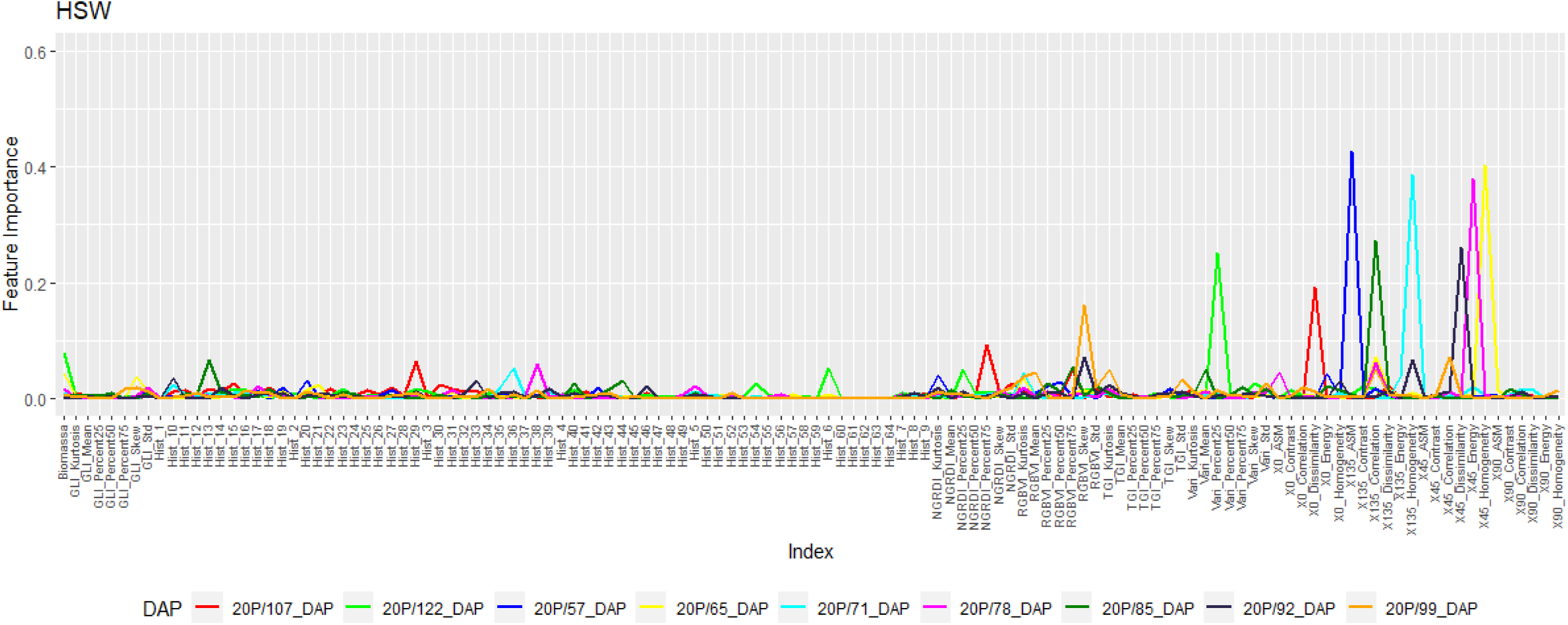
Importance of each predictor variable for predicting HSW in each flight where the predictive ability was greater than 0.60. The importance was calculated based on decision tree analysis, estimating the importance of each image-derived feature using the Gini index.

**Supplementary Fig. 8.**
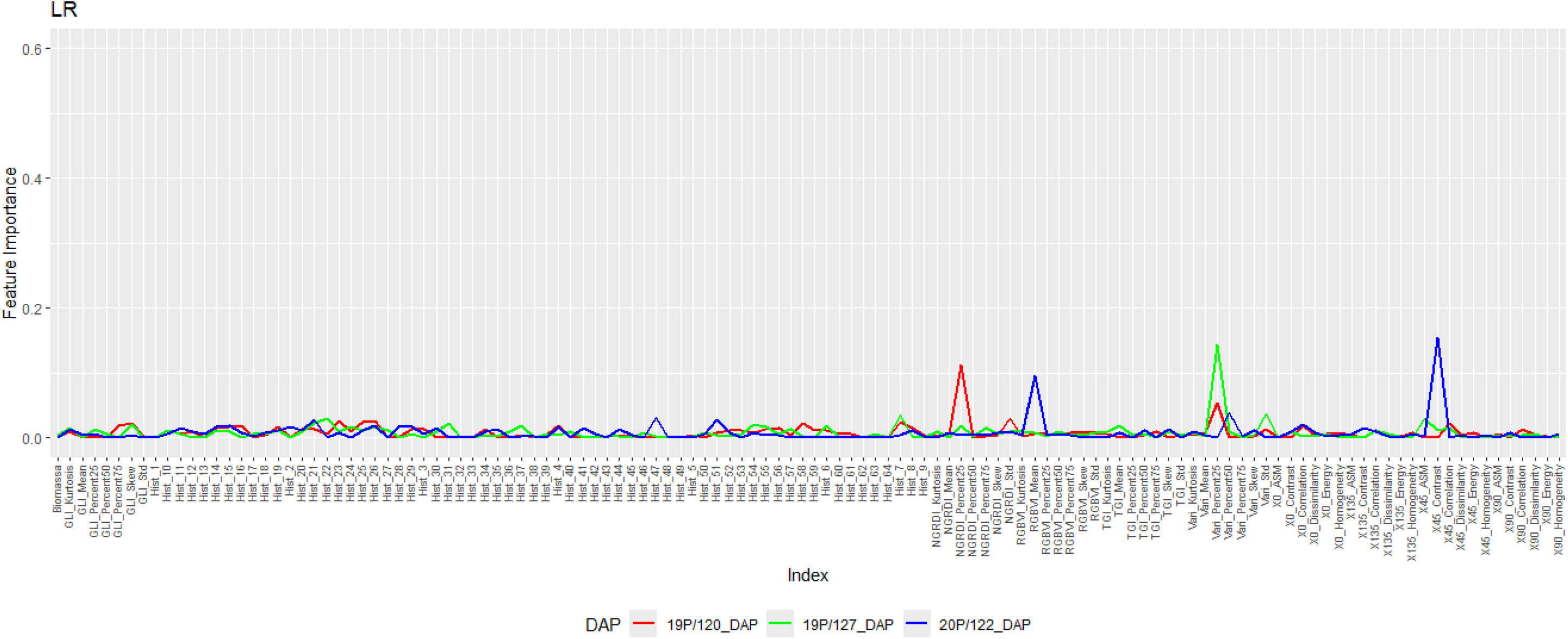
Importance of each predictor variable for predicting LR on flights in which the predictive ability was better. The importance was calculated based on decision tree analysis, estimating the importance of each image-derived feature using the Gini index.

**Supplementary Fig. 9.**
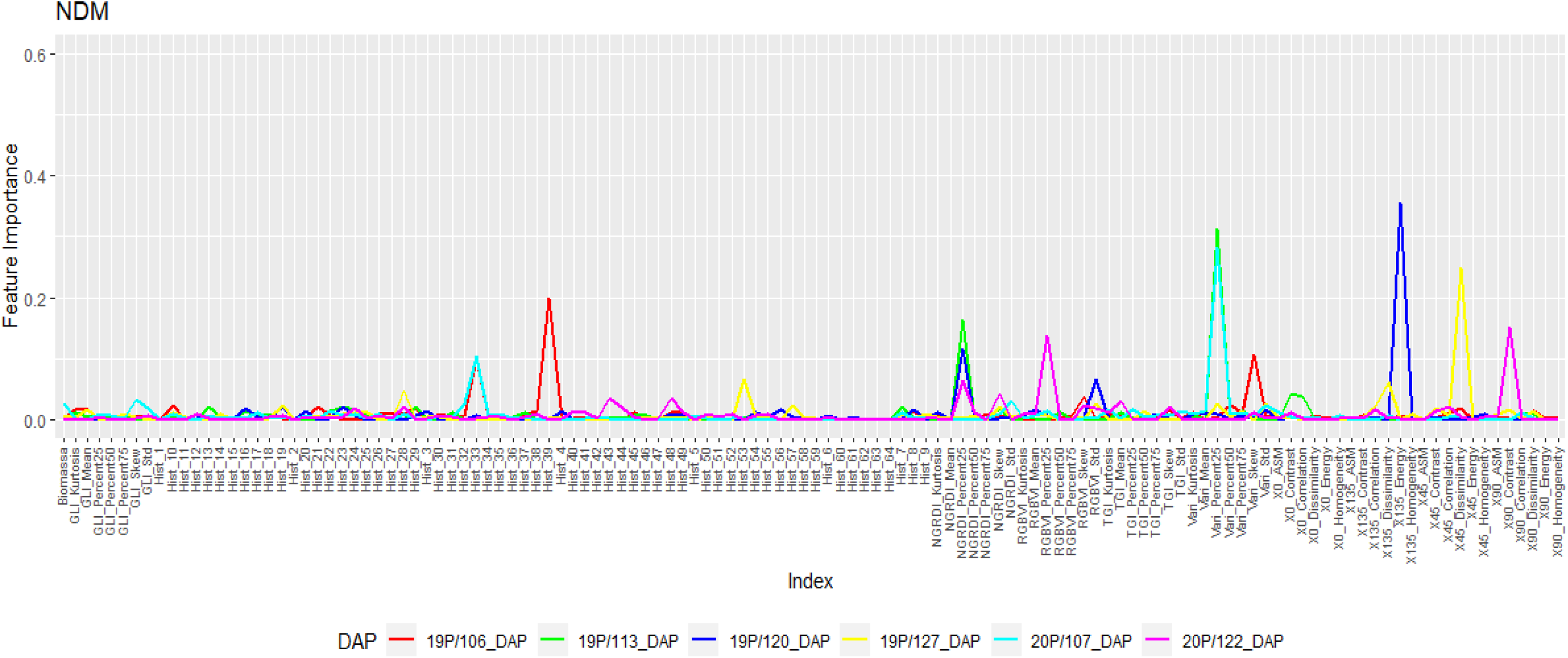
Importance of each predictor variable for predicting NDM in each flight where the predictive ability was greater than 0.60. The importance was calculated based on decision tree analysis, estimating the importance of each image-derived feature using the Gini index.

**Supplementary Fig. 10.**
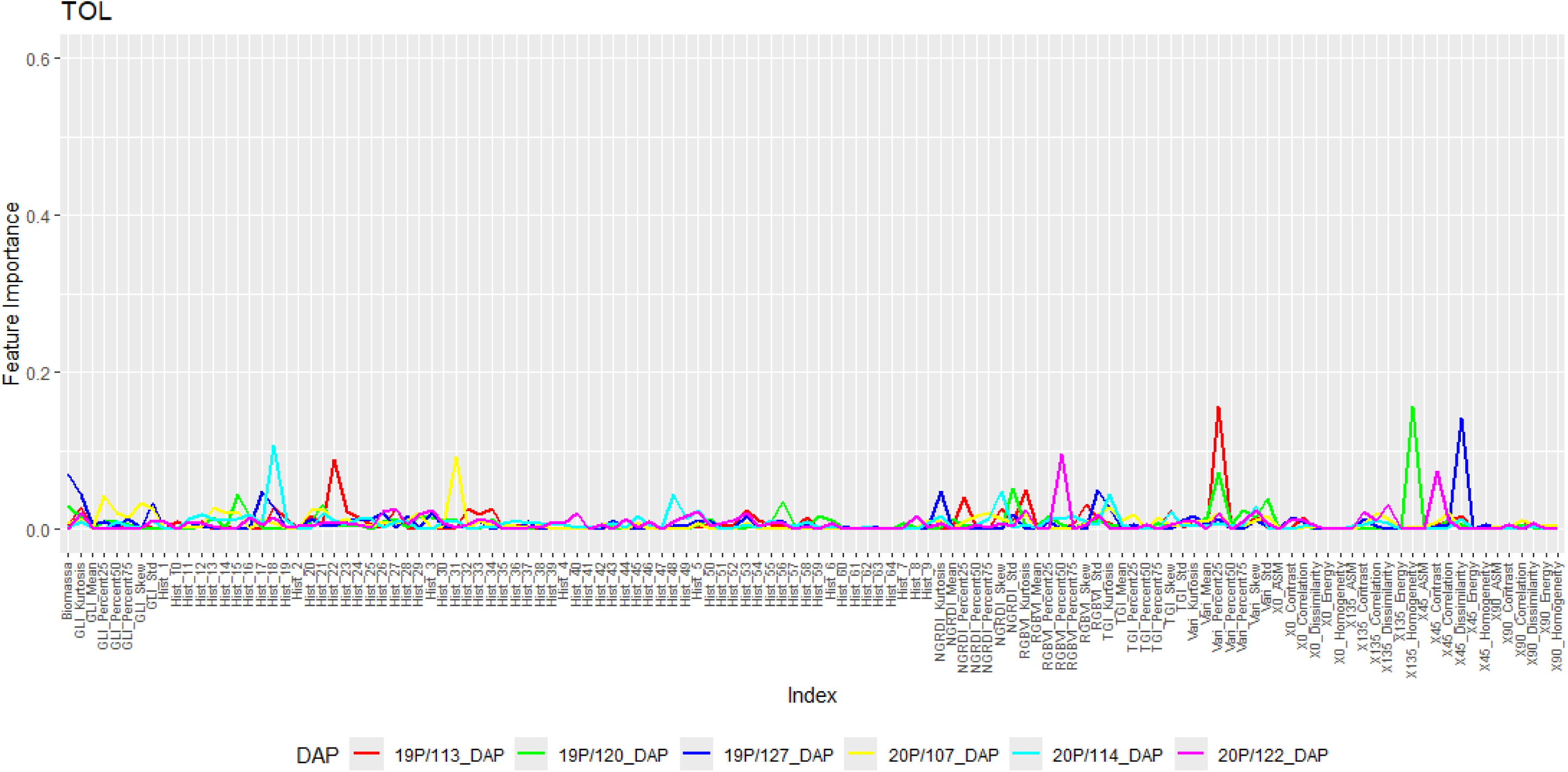
Importance of each predictor variable for predicting NDM on flights in which the predictive ability was better. The importance was calculated based on decision tree analysis, estimating the importance of each image-derived feature using the Gini index.

